# Conceptualising and quantifying phenological plasticity through latent responses to seasonal environmental progression

**DOI:** 10.64898/2026.10.02.755607

**Authors:** Victorine George, Emmanuelle Cam, Jane M. Reid, Paul Acker

## Abstract

Quantifying how individuals adjust the seasonal timings of life-history events (i.e. their phenology) in response to seasonal environmental conditions is fundamental to predicting impacts of changing seasonality. However, the current standard approach of inferring phenological responses by directly analysing observed event dates relies on foundational assumptions that could be invalidated by changing seasonality, and excludes individuals that do not express focal events, potentially generating misleading inferences. To overcome these limitations, we devised a Phenological Threshold Model (PTM) that conceptualizes phenological events as switches between discrete life-history states and primarily estimates variation in underlying latent dynamic responses to seasonal progression, from which event dates secondarily arise. We demonstrate that the PTM can accurately retrieve simulated parameter values, including given incomplete seasonal field observations. Application to multi-year data on annual breeding in black-legged kittiwakes (*Rissa tridactyla*) demonstrated substantial among-individual variation in latent responses to seasonally varying environmental drivers, and associated daily probabilities that individuals will initiate breeding, revealing how annual distributions of breeding dates and proportions of non-breeders jointly arise. We thereby formalise a new conceptual and statistical framework for quantifying the potential for, and constraints on, rapid plastic and micro-evolutionary changes in phenology in changing seasonal climatic and environmental regimes.

## Introduction

Climate change is rapidly modifying seasonal environments, altering the timings and variabilities of seasonal progressions (Hernández-Carrasco et al., 2025; IPBES, 2019). Such changes can disrupt seasonal synchronisation between key life-history events (e.g., flowering, breeding, migration) and their most suitable environments (Burthe et al., 2012; Durant et al., 2007; Parmesan, 2006; Simmonds et al., 2020; Visser et al., 2012). Resulting phenological mismatches can decrease individual survival probability and/or breeding success, potentially reducing population viability (Miller-Rushing et al., 2010; Peng et al., 2025; Radchuk et al., 2019; Visser and Gienapp, 2019). These outcomes substantially depend on the degree to which individuals of focal species can appropriately adjust their seasonal life-history timings through phenological plasticity and/or micro-evolution (Ashander et al., 2016; Bonamour et al., 2019; Charmantier et al., 2008; Chevin et al., 2010; Whelan et al., 2022). Yet, while these concepts are well established, adequate inferences on future phenotypic, evolutionary and demographic responses to changing seasonality still require better understanding of the form, magnitude and basis of individual-level and population-wide variation in phenological responses (Chmura et al., 2019; Donohue et al., 2015; Kharouba and Wolkovich, 2020).

One current standard approach to identifying environmental drivers of phenology, and dissecting individual responses, involves analysing observed annual dates of individuals’ life-history events (e.g., flowering, breeding, migration) as the primary biological trait on which environmental and individual effects directly act. The environment is commonly reductively envisaged as annual ‘cues’, defined as mean values of one or more environmental covariates through fixed time windows preceding the seasonal life-history events (e.g. Bonamour et al., 2019; Gienapp et al., 2013; Whelan et al., 2022). Relevant windows are identified as the time period when among-year variation in the mean environmental value and the mean phenological date are most strongly correlated (Simmonds et al., 2019; Van De Pol et al., 2016). Yet, this conceptualisation effectively assumes that organisms are sensitive to their environment only during a single seasonal window, which is identical for all individuals. This is unlikely, and often evidently untrue (Ball, 1993; Dawson et al., 2001; Gienapp et al., 2005). It also assumes that the ‘cue’ window is fixed, thereby effectively grounding predictions of phenological responses to climate change in a substantially incompatible assumption that the seasonal timing of underlying biological processes will not change (Bonamour et al., 2019; Dawson et al., 2001; Gienapp et al., 2013; Simmonds et al., 2017). Furthermore, directly analyzing observed dates necessarily excludes individuals that do not express a date in a particular year (e.g., do not flower, breed or migrate). Such instances can be commonplace, for example in species where individuals sometimes skip breeding or fail to mate (Cam et al., 1998; Jenouvrier et al., 2003; Lee et al., 2017), and in partially migratory populations (Chapman et al., 2011; Reid et al., 2018). Yet, such individuals are still part of the full population capacity for phenological variation, and for evolution of phenology and phenological plasticity. Ignoring these individuals could be highly problematic as they are likely ‘missing not at random’ with respect to expected phenology. Their omission could therefore bias estimates of trait variation and any subsequent inferences on underlying genetic variation, natural selection and resulting evolution (e.g. Hadfield, 2008; Mojica and Kelly, 2010). Frequently reported discrepancies between observed and predicted phenological responses (Keogan et al., 2018; Pujol et al., 2018) may therefore partly arise because direct date-based analyses misconceptualise, or at least over-simplify, phenology and consequently do not adequately capture realistic forms of individual response or associated constraints (Chmura et al., 2019; Gienapp et al., 2005, 2013; Kharouba and Wolkovich, 2020; Macphie et al., 2026; Reid and Acker, 2022a).

Overcoming these limitations requires developing alternative conceptual and analytical frameworks that more explicitly envisage and test mechanistic hypotheses linking seasonally-varying environmental conditions to individual phenological outcomes (including failure to express the focal life-history event; Chmura et al., 2019; Donohue et al., 2015; Inouye et al., 2019; Kharouba and Wolkovich, 2020). This can be achieved using the ‘threshold trait’ framework, which envisages that underlying individual characteristics and seasonally dynamic environmental conditions combine to generate a switch between discrete life-history states that define phenology (e.g., not yet flowering, breeding, or migrated to flowering, breeding or migrated, Figure 1A; Reid and Acker, 2022). The state switch takes place when a continuously distributed underlying latent variable (termed ‘liability’; Falconer, 1965) exceeds a certain threshold. The switch can occur from day to day within each year, generating an observable switch date (Figure 1). Among-individual variation in date then results from among-individual variation in liability-scale intercepts and/or slopes on one or multiple environmental variables that change as each season progresses. This could include variables that are effectively deterministic or stochastic functions of time (e.g. photoperiod, temperature), which could act immediately or cumulatively over some within-season timeframe (Arnold et al., 2019; Ball and Balthazart, 2003; Chuine and Régnière, 2017; Dawson et al., 2001). Correspondingly, individuals that do not express a date within a particular year are those whose liabilities do not respond sufficiently to reach the threshold to switch state (Figure 1A-C, Reid and Acker, 2022a, b).

**Figure 1.**
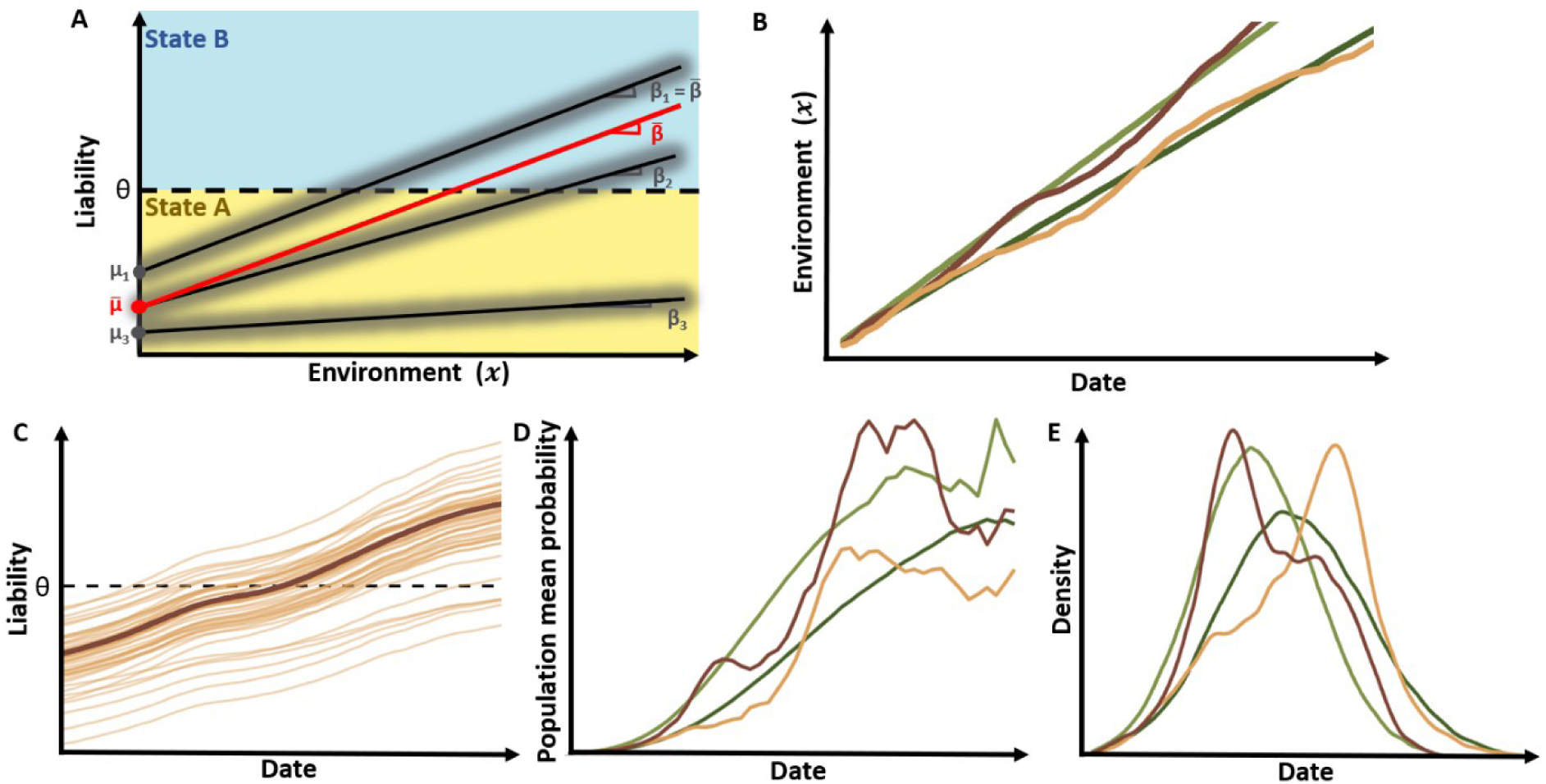
Conceptual framework of the Phenological Threshold Model (PTM). A) Latent Reaction Norms (LRNs) for three illustrative individuals (black lines) and the population mean (red line) given some seasonal environmental variable, x. Individuals switch between two alternative phenotypic states (A, yellow and B, blue) when their liabilities (l, black lines) cross the threshold (θ, dashed line). The LRNs are defined by the intercept µ (giving the initial distance from the threshold), and the slope β (representing the sensitivity to x), both of which can vary among individuals. The LRNs are depicted as linear functions of x but could in principle take other (non-linear) shapes. Surrounding shading represents additional residual variation in the liability response. B) Examples of seasonal trajectories of two hypothetical environmental variables varying between two years (dark and light for years 1 and 2), which monotonically increase with date with differing gradients, either linearly (green) or non-linearly (brown). C) Examples of individuals’ liability responses to the non-linear environmental variable through the season in year 1, where the dark line denotes the population mean and the lighter lines are individuals’ responses. For visual clarity, daily residual variations are not shown. D) Population mean seasonal trajectories of the daily probability that an individual realises the life-history event and thus switches state (conditional on not having switched yet), Ψ^-^_d,y_, and E) resulting density distributions of observable phenological dates given the environmental variables depicted in panel B.

By focusing on environmentally-induced processes underlying the potential phenotypic switch, rather than solely on realised switches, the threshold trait framework facilitates explicit formulation and test of hypotheses concerning biological responses to changing seasonality, and hence prediction of phenological responses to future environmental regimes. Specifically, by considering latent reaction norms (Figure 1C), it can predict individual probabilities of threshold crossing (i.e. state switching) on each day (Figure 1D) and hence predict resulting distributions of phenological dates (Figure 1E) for any set of observed or projected environmental drivers (Figure 1B) without invoking fixed preceding cue windows or excluding non-crossing individuals. It could also identify current hard constraints, such as a lack of evident among-individual variation in responses to components of seasonality, which would imply limited potential for rapid evolutionary change in phenology. Moreover, this framework allows such analyses given driving seasonal environmental variables that do not vary among years (e.g. photoperiod), variables that do vary among years (e.g., temperature) and their potential interactions (e.g. photoperiod-by-temperature), which is not possible using a fixed cue window approach. Overall, the threshold trait approach could thereby provide multiple new insights that enhance previous inferences resulting from traditional analyses of observed dates.

While the threshold trait approach and its core concepts of liabilities and latent reaction norms (Figure 1) might seem complex and abstract, it in fact invokes exactly the same construction as is widely used to analyse any binary response variable (e.g. logistic regression or binary generalized linear (mixed) models (GLMMs), Acker and Reid, 2025). The extension to study phenology simply requires envisaging a temporal (e.g. daily) sequence of individual-level binary outcomes manifested throughout a season, which can be induced by underlying seasonally varying covariates. However, although seasonally longitudinal approaches to studying population-level phenology were previously established, the shift from date-centric to binary trait analyses has not yet been explicitly formulated or enacted in the context of quantifying among-individual variation in reaction norms of plasticity in evolutionary ecology (Donohue et al., 2015; Gienapp et al., 2005; Reid and Acker, 2022a). It is consequently unclear how effectively key parameters can be estimated from field data, including the common scenario where individuals’ states are not observed every day, and resulting conceptual and quantitative insights have not been exemplified.

Accordingly, we devised a general ‘phenological threshold model’ (PTM) capable of statistically estimating latent reaction norm parameters describing liability responses to environmental drivers of seasonal life-history state switching, and verified its efficacy given simulated data. Further, we show how embedding a Hidden Markov (HM) model within the PTM can remove biases stemming from intermittent rather than systematic daily observations of individuals’ states, thereby handling observation gaps that are commonplace in field studies. To illustrate resulting biological insights, we fitted a HM-PTM to annual nest completion data, demarcating initiation of breeding, in individually-marked kittiwakes (*Rissa tridactyla*). We demonstrate substantial among-individual variation in liability-scale responses to illustrative seasonal environmental variables, and reveal how annual distributions of nest completion dates and frequencies of non-breeding jointly emerge. We thereby demonstrate how reconceptualising phenology as a threshold trait can provide novel insights into the degree to which climate-threatened populations harbour potential to rapidly respond to changing seasonal environments, affecting both the occurrence and timing of key life-history events.

### Phenological Threshold Model

#### Model formulation

To formalise and demonstrate the PTM, we envisage a phenological trait comprising two alternative life-history states, expressed in a population where known individuals are monitored as the season progresses in each of several years. Mostly simply, we conceptualise a liability-scale latent reaction norm (LRN), where each individual *i* has an intercept, *μ_i_*, and a slope, *β_x_*_,*i*_, defining its liability response to an environmental variable *x*, where *x* has value *x_d_*_,*y*_ on day *d* in year *y* (Figure 1A). This construction is general, and handles environmental variables that do or do not vary in *d* and/or *y*. It is readily extended to be multivariate, invoking multiple environmental variables. Given that seasonal progressions of some variables (e.g. temperature) vary among years (e.g. Figure 1B), seasonal trajectories of individual liabilities can vary among years even if *μ_i_* and *β_x_*_,*i*_ are constant (Figure 1C). Quantifying population-wide variation in phenological plasticity then requires quantifying the population mean liability intercept μ^-^ and slope β^-^*_x_* defining the average response to each focal environmental covariate, and quantifying the individual deviations *δ_μ_*_,*i*_ and *δ_βx_*_,*i*_ in intercept and slope summarised as the among-individual variances (σ^2^_μ_ and σ^2^_βx_; Figure 1A).

To achieve this using typically-collected field data, for each individual *i* in each year *y* we envisage the observed phenological data as a state sequence, *H_i_*_,*y*_. Here, on each day *d* throughout a study season of duration D days within *y*, *i* is observed in state *S_i_*_,*d*,*y*_ which is either ‘A’ or ‘B’, representing the pre-and post-event phenotypes (Figure 2A). The annual observations are assumed to start before any individual expresses the state-switch event, and finish after all individuals who will switch within year *y* have done so. Considering for current purposes that the phenological trait is irreversible within years (e.g. individuals do not revert to pre-breeding or pre-migration after breeding or migrating, Supplemental material B), all individuals start the season in state A, and on each subsequent day can either remain in state A or switch to state B (e.g. Figure 2A). The probability that *i* switches on day *d* (conditional on not having switched previously) is then *Ψ_i_*_,*d*,*y*_ = *P*(*S_i_*_,*d*,*y*_ = *B*|*S_i_*_,*d*−1,*y*_ = *A*).

**Figure 2.**
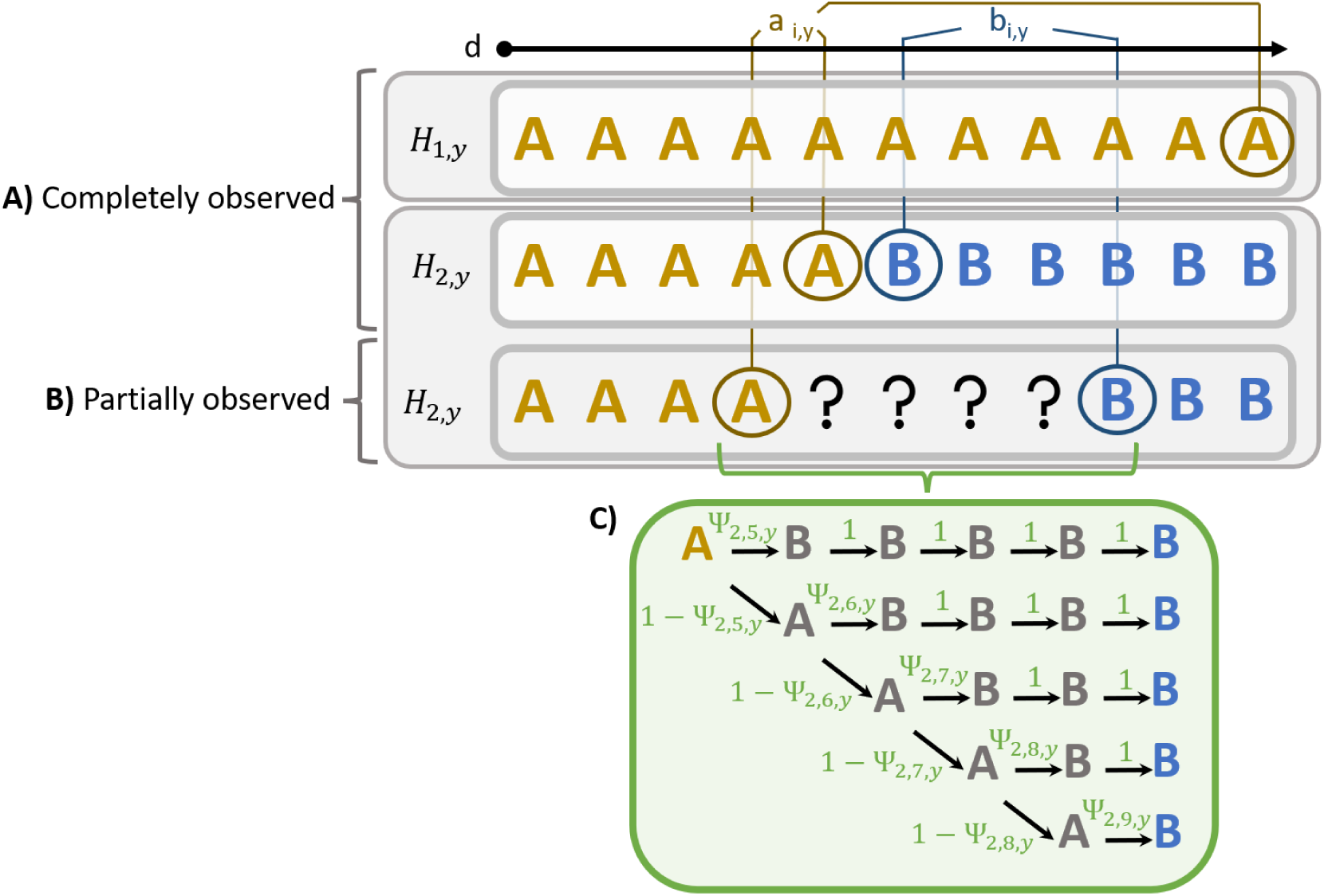
Conceptual representation of the Hidden Markov Phenological Threshold Model (HM-PTM) for an irreversible trait with alternative phenotypic states A and B. A) Complete observed state sequences (H_i,y_) of two individuals (i=1 and i=2) through 11 days. Individual 1 remains in state A, while individual 2 switches state between days 5 and 6. B) Incomplete observed state sequence of individual 2, given no field observation from day 5 to 8 (question marks). The phenotypic state S_i,d,y_ is therefore unknown from the day after i was last observed in state A, a_i,y_, until the first day it was observed in state B, b_i,y_. C) Probability tree illustrating the Hidden Markov model for the missing observations in H_2,y_. Here, **Ψ**_2,d,y_ is the probability that individual 2 switched to B on day d in year y, conditional on not having switched previously within that year. The resulting probability that individual 2 switched on any particular day after a_2,y_, χ_2,a2,y,y_, is the sum of all path probabilities in the tree (where each path probability is the product of the probabilities of the successive branches defining the path; Supplemental material D). This formulation assumes that, conditional on an observation being made, states A or B are correctly recorded without error.

We then formulate a PTM to estimate the LRN parameters that generate *S_i_*_,*d*,*y*_, and the underlying *Ψ_i_*_,*d*,*y*_, given any focal set of *x_d_*_,*y*_ values. This is enacted by fitting a longitudinal binary GLMM to the observed *H_i_*_,*y*_ data, where *Ψ_i_*_,*d*,*y*_ is a function of the LRN parameters (i.e. μ^-^, β^-^*_x_*, σ^2^_μ_ and σ^2^_βx_). This is achieved using a logit link (or other appropriate link, e.g. probit or cloglog), defining a standard transformation between the dichotomous phenotypic scale and the latent liability scale (Equation 1, Supplemental material C; Acker and Reid, 2025):

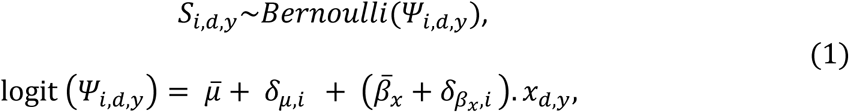

where the individual deviations *δ_μ_*_,*i*_ ∼N(0, σ^2^_μ_) and *δ_βx_*_,*I*_ ∼N(0, σ^2^_μx_) (i.e. normally distributed random effects). This standardized model formulates the LRN parameters on a logit scale, centred on the threshold with unit √3/π times the standard deviation of the latent daily residuals (Acker and Reid, 2025).

When *S_i_*_,*d*,*y*_ is observed for each day in each focal year with no missing data or error (Figure 2A), the GLMM is effectively fitted to all successive binary observations up to the first day when *i* is observed in state B, *b_i_*_,*y*_, or the last day it is observed in state A, *a_i_*_,*y*_, if it never switches state. The probability of the observed sequence *H_i_*_,*y*_, *P*(*H_i_*_,*y*_), defining the likelihood for *H_i_*_,*y*_, is then:

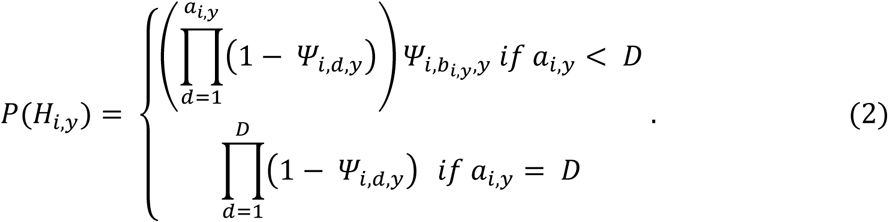

Assuming independence across individuals, the full model likelihood is given by the product of all *P*(*H_i_*_,*y*_).

However, in reality, *H_i_*_,*y*_ will often include gaps with unknown *S_i_*_,*d*,*y*_ (Figure 2B), for example if field visits are not undertaken every day or if field conditions prevent some planned observations. The actual days when individuals switched from state A to state B are then unknown. In such circumstances, switching dates may sometimes be reasonably back-casted from subsequent observations (e.g. inferring first laying date given observed subsequent clutch size by assuming one egg laid per day; Evans et al., 2020; Supplemental material B). However, in other cases (e.g. flowering or migration dates), such back-casting cannot necessarily be implemented. Fitting the basic PTM (Equation 2) to incomplete *H_i_*_,*y*_ data could then yield substantially biased estimates of LRN parameters, because the switch could have occurred earlier (but not later) than *b_i_*_,*y*_. Hence, to appropriately handle missing data, we embedded a Hidden Markov (HM) model within the PTM, formulating the probability *χ_i_*_,*a*_*_i_*_,*y*,*y*_ that *i* switches on any particular day after *a_i_*_,*y*_ and no later than *b_i_*_,*y*_ (if *b_i_*_,*y*_ exists; Figure 2C). Then, *P*(*H_i_*_,*y*_) can be expressed as:

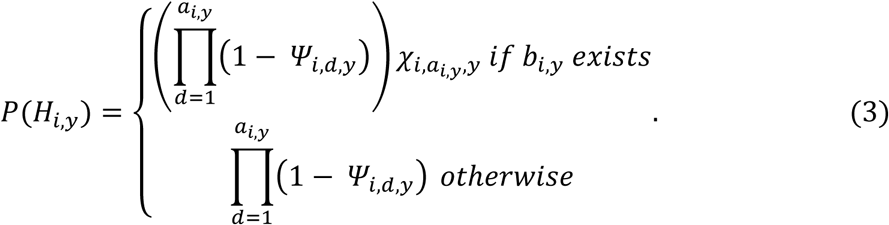

More generally given the specified model, we can define the probability that *i* switches on any particular day after any day *t* and no later than *b_i_*_,*y*_ (conditional on not having switched yet on day *t*), *χ_i_*_,*t*,*y*_ as:

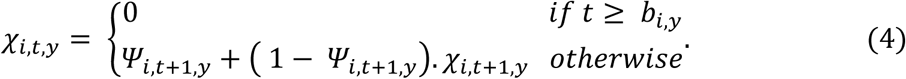

This formulation makes *χ_i_*_,*a*_*_i_*_,*y*,*y*_ readily calculable using a backward recursion from *t* = *b_i_*_,*y*_ (Supplemental material D). Equation 3 also handles cases where *i* is never observed in state B, including due to incomplete late season observations, leaving uncertainty on whether it ever switched or not.

#### Model evaluation

We conducted simulations to verify that the HM-PTM accurately retrieves underlying LRN parameters from observed *H_i_*_,*y*_. We first simulated two sets of an environmental variable *x*, with linear or non-linear seasonal trajectories through an annual study period of D=50 days within each of six years (e.g. Figure 1B). The non-linear trajectories were generated as a first-order autoregressive process (Supplemental material E), emulating a stochastically progressing seasonal environment (e.g. rolling mean temperature or cumulative growing degree days). We then generated each individual’s annual liability trajectory given one or the other covariate and specified values of the four population-level LRN parameters (μ^-^, β^-^*_x_*, σ^2^_μ_ and σ^2^_μx_, Supplemental material E). To evaluate a basic model, then verify that both σ^2^_μ_ and σ^2^_μx_ were identifiable, we created two sets of parameter values with zero or non-zero σ^2^_μx_. Values were chosen to ensure that all individuals started with liabilities below the threshold and most, but not necessarily all, crossed the threshold before D. Phenological state sequences were generated by identifying the first day when each individual’s liability crossed the threshold, before which *S_i_*_,*d*,*y*_ is A and from which *S_i_*_,*d*,*y*_ becomes and remains B.

For each covariate and set of parameter values, we generated two datasets. The first envisaged a fully balanced design, with 1000 individuals that were all observed in all six simulated years. The second envisaged an unbalanced design, with 3706 individuals that were present in differing number of years within the possible six, with most individuals present only once (representing a more realistic situation where individuals recruit and die in different years). Both datasets contained 6000 *H_i_*_,*y*_ in total. We then either retained individuals’ full state sequences (yielding complete *H_i_*_,*y*_) or degraded them by generating missing observations envisaging a situation where field data are collected every sixth day (yielding incomplete *H_i_*_,*y*_ with gaps). This generated sixteen simulated datasets spanning linear or stochastic seasonal environmental progression, with or without among-individual variation in LRN slopes, balanced or unbalanced, and complete or gappy (Supplemental material E).

We fitted the basic PTM to all complete datasets and to the balanced gappy datasets, and the HM-PTM to all gappy datasets, using Bayesian inference coded in Stan called from R (R core team 2026) with package rstan (Carpenter et al., 2017). To evaluate model performance, we compared the posterior means and 95% credible intervals (95%CRI) of μ^-^, β^-^*_x_*, σ^2^_μ_, and σ^2^_μx_ to the simulated values. Codes for simulations and analyses are available in (Anonymous, 2026).

These analyses showed that the PTM can accurately retrieve the simulated LRN parameters given complete *H_i_*_,*y*_ data, with linear or stochastic environmental covariates and balanced or unbalanced designs. Here, the 95%CRI for μ^-^, β^-^*_x_*, σ^2^_μ_ and σ^2^_μx_ are narrow and encompass the simulated values with no systematic bias (Figure 3). However, as expected, degraded gappy datasets induce substantial biases (Figure 3). Specifically, μ^-^ and β^-^*_x_* show severe downward and upward bias respectively, implying underestimation of mean liability at the defined seasonal starting point, and overestimation of mean sensitivity to the seasonal environmental progression. The among-individual variances are also upwardly biased, implying overestimation of the within-population variation in both initial liability and seasonal sensitivity. These important biases, which would generate misleading biological interpretations, are successfully corrected by fitting the HM-PTM (Figure 3).

**Figure 3.**
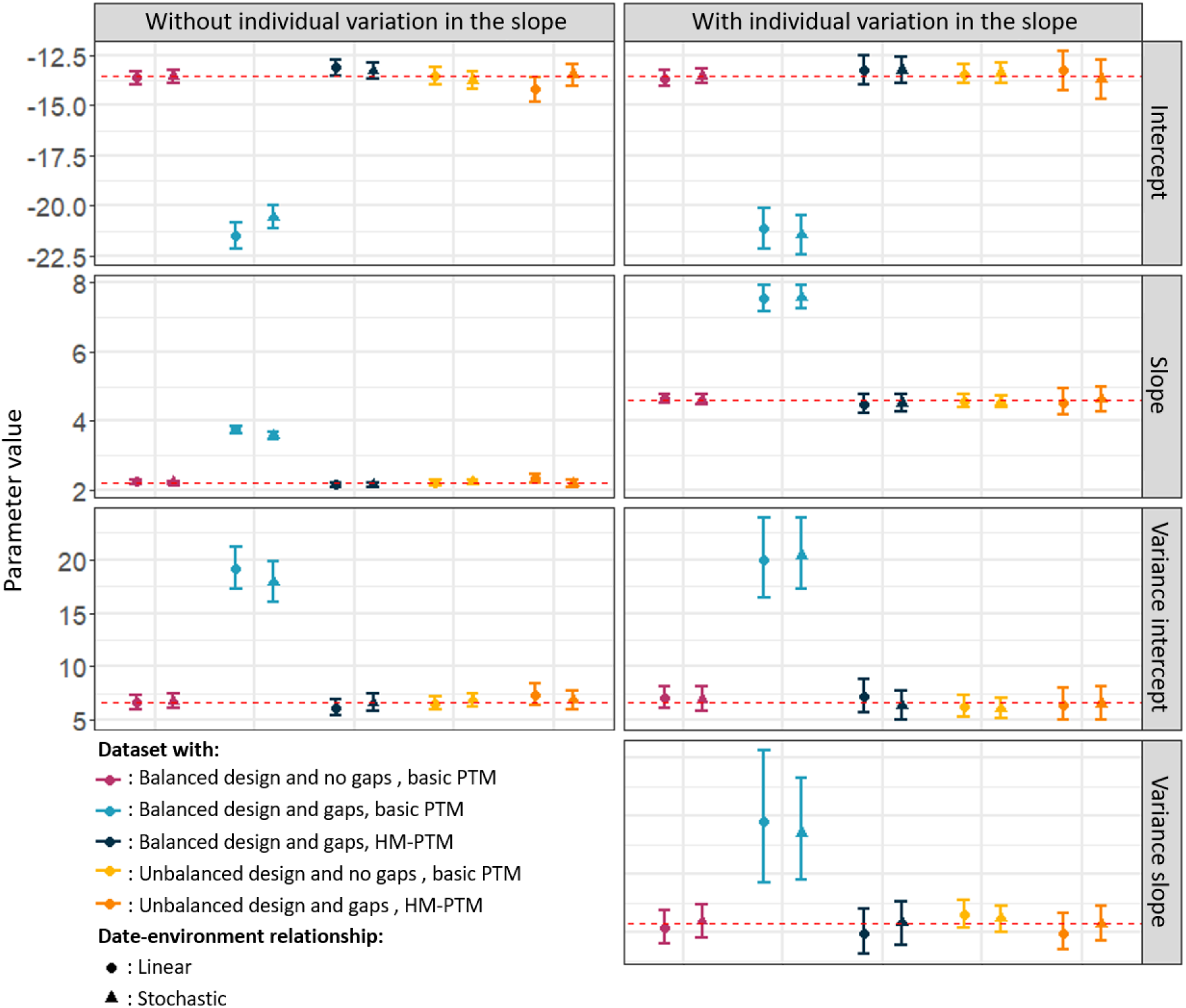
Simulated (red dashed lines) and estimated parameter values and 95%CRI (points and associated bars) for each simulated latent reaction norm (LRN) dataset. Rows depict the four LRN parameters (top to bottom: population mean intercept, µ̅, and slope, β̅_x_, and among-individual variance in the intercept, σ^2^_μ_, and slope σ^2^_μx_). Left and right columns show simulations without and with among-individual variation in the slope, respectively (i.e., zero and non-zero σ^2^). Colours and symbols denote different simulated datasets analysed with the basic phenological threshold model (PTM) or with the additional Hidden Markov component (HM-PTM, details in bottom-left legend).

#### Model implications and interpretations

The demonstration that the (HM-)PTM can in principle accurately estimate key LRN parameters defining the relationship between liability and seasonal environmental progression from (incomplete) state-sequence data (Figures 1-3) opens the door to operationalising a reconceptualization of phenological dynamics, yielding three notable opportunities. First, it could allow tests of mechanistic hypotheses on effects of fully predictable year-invariant seasonal drivers of phenology (e.g. photoperiod) alongside stochastic seasonal drivers whose trajectories vary among years (e.g. absolute and/or cumulative weather variables). This would be achieved by testing which LRN slopes and among-individual variances differ from zero. Among-individual variances could reflect previous developmental plasticity, and/or genetic variation (potentially allowing rapid evolution of phenology). Such effects involving year-invariant drivers cannot be quantified through standard date and cue window analyses (which require among-year variation; Chmura et al., 2019; Dawson, 2015; Hahn et al., 2025).

Second, the (HM-)PTM allows computation of individual-level and population-level daily probabilities of expressing the focal life-history switch (conditional on not having previously done so) in each year (*Ψ_i_*_,*d,y*_ and ψ^-^*_d_*_,*y*_ respectively, Figure 1D) given all modelled individual and environmental effects. These probabilities effectively stem from the means and among-individual variances in the LRN intercept (μ^-^ and σ^2^_μ_) and slopes (β^-^*_x_* and σ^2^_μx_) on seasonal environmental variables, given the defined distribution of the daily residual variance in liability (Supplemental material C; Acker and Reid, 2025). By summarising the net daily joint action of the underlying environmental drivers on the expected state switch, *Ψ_i_*_,*d,y*_ and ψ^-^*_d_*_,*y*_ identify the non-linear trajectory of the focal life-history trait response through the season.

This is central to understanding how phenology emerges from seasonal environmental progression, and allows testing hypotheses involving phenological dynamics (i.e. when, and why, seasonal trajectories of switching rates begin, accelerate, slow down, stop, or reverse).

Accordingly, third, calculating ψ^-^*_d_*_,*y*_ (Figure 1D) given a set of estimated LRN parameters (Figure 1A) and environmental values (Figure 1B) enables jointly predicting annual distributions of realized phenological dates (Figure 1E) and proportions of individuals that do not express the focal life-history switch in each year. The (HM-)PTM thereby reveals how differing annual distributions of phenological dates could emerge from responses to among-year variation in within-year seasonal environmental progression, and how differing annual frequencies of non-expression can simultaneously arise from the same biological effects. Given incomplete field observations (gappy *H_i_*_,*y*_), predicted dates and non-expression frequencies inferred from the HM-PTM could be more accurate than the raw observed dates and frequencies, since the bias stemming from observation failure will be substantially reduced (Supplemental material J). This could facilitate testing hypotheses concerning dates, including testing for selection occurring through phenological mismatches.

### Application of the Phenological Threshold Model

#### Study system and phenological trait

To demonstrate the application and utility of the threshold trait framework, we fitted an HM-PTM to a multi-year dataset on annual breeding times from black-legged kittiwakes breeding in Cap Sizun (48°03’N, 4°39’W), France (Acker et al., 2022; Cam et al., 1998). Since 1979, large proportions of chicks were individually colour-ringed, allowing subsequent monitoring of numerous individuals that locally recruited. Since 1883, comprehensive surveys of all previous and actual nesting sites were conducted during January-July, typically every 1-4 days during 1983-2003 and every 5-12 days subsequently (Supplemental material F). Systematic resighting effort yielded virtually complete annual detection of recruited adults (probability≍0.99) and identification of all ringed nest owners (Acker et al., 2022; Cam et al., 1998). This monitoring enabled recording timings of individuals’ life-history events within and among-years, spanning diverse environmental conditions (Supplemental material F; Acker et al., 2022; Cam et al., 1998). For current illustrative purposes, we focus on the trait of nest completion, which is prerequisite for reproduction and a classical criterion used for distinguishing breeders from non-breeders in kittiwakes (Supplemental material F; Cullen, 1957; Danchin, 1987; Maunder and Threlfall, 1972).

#### Seasonal covariates

Fitting a (HM-)PTM requires appropriate covariates that drive phenological progression within years, and variation among years. For current illustrative purposes, we defined two photoperiod-related variables: daylength (φ*_d_*, number of hours between sunrise and sunset on day *d*), and the difference in daylength between consecutive days (Δφ*_d_* = φ*_d_* − φ*_d_*_−1_, in hours). These two variables capture known processes of reproductive photoinduction, regulating the onset of breeding activities, and photoinhibition, leading to reproductive cessation (see Discussion, Supplemental material G; Ball and Balthazart, 2003; Dawson, 2015; Dawson et al., 2001; Watanabe et al., 2007). Together, they generate appropriate limits on the annual reproductive season (Supplemental material G).

Yet, photoperiod cannot explain among-year variation in reproductive timing, which commonly varies with stochastic environmental variables that shape seasonal biological progression (e.g., temperature, rainfall; Ball and Balthazart, 2003; Bonamour et al., 2019; Maunder and Threlfall, 1972; Moe et al., 2009). For current illustrative purposes, we defined a temperature anomaly variable (*T*, in °C) that represents within- and among-year variation in temperature while avoiding collinearity with φ*_d_* and Δφ*_d_*. We computed *T_d_*_,*y*_ for each day *d* in each year *y* as the deviation of the mean of the daily air temperature averaged over 20 preceding days (up to and including each focal day) from the multi-year grand mean for that day. When applied for each day in each year this procedure effectively generates a rolling mean temperature anomaly, with the 20-day interval chosen to capture expected biological and physical effects (e.g. on sea surface temperature) which arise across longer time periods than single days. Daily mean temperature measured at the meteorological station closest to the focal kittiwake colonies were extracted from the Météo-France online database (Météo-France, 2024, Supplemental material G). Finally, φ*_d_* and Δφ*_d_* were standardised (i.e. mean-centred and divided by the sample standard deviation) and *T_d_*_,*y*_ was scaled (i.e. divided by the sample standard deviation) to facilitate computation and interpretations.

#### Life-history state dataset

To generate observed state sequences *H_i_*_,*y*_ for each individual *i* in year *y* we assigned state A on all days up to and including the last day its nest was observed as incomplete, state B the day its nest was first observed to be complete, or NA when its nest was not observed on days after the last A and preceding the first B, and after the first B until the defined season end D (Supplemental material F). Individuals that were first observed with a completed nest were assigned state A on *d*=1, then NA up to the day before the first observation in state B. All *H_i_*_,*y*_ were defined over D=125 days spanning ordinal dates 76–200 (March 18th to July 19th).

Ordinal date 76 was specified because it is 14 days before the earliest ever observed completed nest (March 31^st^, ordinal date 90), thereby encompassing the full range of observed phenology plus the maximum potential for observation failure (Supplemental material F). Ordinal date 200 is the latest ever recorded completed nest. All ringed adults that bred at least once within the study area were included in each year in which they were resighted, irrespective of whether they completed a nest that year. We used data from 1985 to 2022, instead of 1983, because the average age at first breeding is 4 years old (Danchin and Cam, 2002; Supplemental material F). Kittiwake data for 2014-2019 were not used because temperature data were missing, precluding calculation of *T_d_*_,*y*_. Resulting dataset comprises a total of 32 years and 13,113 *H_i_*_,*y*_ from 3,209 recruited adults.

#### Statistical analyses and inferences

We fitted a HM-PTM to estimate the LRN parameters assumed to underlie the *H_i_*_,*y*_. Specifically, the probability *Ψ_i_*_,*d,y*_ that *i* achieves nest completion on day *d* in year *y* (conditional on not having achieved it yet that year) was specified as:

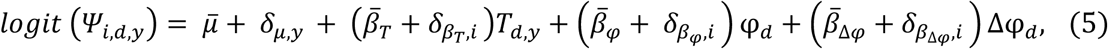

where *δ_μ_*_,*y*_ is an annual deviation from the mean LRN intercept accounting for among-year variation affecting the whole population independently of *T_d_*_,*y*_; β^-^*_T_*,, β^-^*_φ_* and β^-^_Δ*φ*_ are the population mean LRN slopes for the defined covariates *T, φ* and Δ*φ*, and *δ_βT_*_,*i*_, *δ_βφ_*_,*i*_ and *δ_β_* _,*i*_ are the individual deviations in these slopes, distributed such that *δ_β_* _,*i*_∼N(0,σ^2^_μ_) *δ_β_* _,*i*_ ∼N(0, σ^2^_μx_), and *δ_β_* _,*i*_ ∼N(0, σ^2^_μ_). We did not model among-individual variation in the intercept (i.e. no *δ_μ_*_,*i*_) because this is redundant with *δ_βφ_*_,*i*_ and *δ_β_*_Δ*φ,i*_ given that *φ_d_* does not vary among years (Supplemental material I). All modelled among-individual variation therefore concerns sensitivity to the focal environmental covariates.

We performed Bayesian inference using Stan called from R, with weakly informative priors appropriate for the scale of each parameter (Supplemental material H; Anonymous, 2026), and four MCMC chains of 6,500 iterations including 2,000 warm-ups, yielding 18,000 posterior samples. Visual inspection and numerical diagnostics indicated good convergence, with all Ř < 1.01, and effective sample sizes >400 (Supplemental material H; Vehtari et al., 2021).

To predict the population-level daily probabilities of nest completion, ψ^-^*_d_*_,*y*_, the distribution of completion dates and the proportion of non-breeders, *Ω_y_*, in each year, we generated posterior predictions of individual liability trajectories, and resulting state sequences, given the year-specific environmental covariates. This was done for each posterior sample (thereby generating full posterior distributions), by assigning the sampled population and individual parameter values from the HM-PTM (Equation 5) to draw each individual’s phenological state sequence. ψ^-^*_d_*_,*y*_ was then calculated as the proportion of individuals switching to state B on day *d* among those that were in state A on *d*-1. The probability mass function of the annual distribution of nest completion dates was calculated as the proportion of individuals that switched state on each day among all individuals switching state that year. *Ω_y_* was derived as the proportion of individuals that never switched state in the focal year.

Finally, we compared the model-predicted posterior distributions of nest completion dates and *Ω_y_* for each year with the raw observational data. The difference between the two can be interpreted as the bias in estimated phenology as directly inferred from the raw incomplete field observations, alongside any lack of model fit (Supplemental material F and I).

## Results

### Model estimates

As envisaged by construction given that no individuals had already achieved nest completion prior to the analysed annual timeframe, the HM-PTM estimated a negative overall liability-scale intercept μ^-^ (Table 1). There was also substantial among-year variance in the intercept (posterior distribution of σ^2^_μ,y_ peaking clearly away from zero; Table 1; Supplemental material H). This implies that initial (late winter) liabilities are effectively closer to the threshold in some years than others, setting the potential for among-year variation in nest completion timings irrespective of any among-year variation in subsequent seasonal environmental progression. Population mean liability slopes on both defined covariates describing seasonal progression through daylength were non-zero (β^-^*_φ_* and β^-^_Δ*φ*_ ≠ 0, Table 1). As expected, the mean effect of daylength (β^-^*_φ_*) was strongly positive, capturing the increasing probability of nest completion with increasing daylength (i.e. photoinduction). The effect of the change in daylength (β^-^_Δ*φ*_) was also strongly positive, implying a decreasing probability of nest completion with a decreasing difference in daylength between consecutive days, capturing photoinhibition towards midsummer. There was also notable among-individual variation in slopes on both φ*_d_* and Δφ*_d_*: σ^2^_μϕ_ and σ^2^_μΔϕ_ had posterior distributions peaking away from zero (Table 1; Supplemental material H). This implies that individuals vary in their responses to fixed components of seasonal progression.

**Table 1:** Parameter estimates (posterior mean, mode, and median) from the HM-PTM fitted to the kittiwake data, with 95%CRI.

| Parameter | Posterior summary |  |  |  |
| --- | --- | --- | --- | --- |
|  | Mean | Mode | Median | 95%CRI |
| Intercept ($\bar{\mu}$) | -6.760 | -6.750 | -6.760 | [-6.960; -6.560] |
| Slope on T ($\bar{\beta}_T$) | -0.161 | -0.160 | -0.161 | [-0.197; -0.126] |
| Slope on $\varphi$ ($\bar{\beta}_\varphi$) | 5.470 | 5.460 | 5.470 | [5.330; 5.610] |
| Slope on $\Delta\varphi$ ($\bar{\beta}_{\Delta\varphi}$) | 5.710 | 5.680 | 5.710 | [5.490; 5.930] |
| Variance in individual deviations from $\bar{\beta}_T$ ($\sigma_{\beta_T}^2$) | 0.005 | 0.000 | 0.003 | [0.000; 0.016] |
| Variance in individual deviations from $\bar{\beta}_\varphi$ ($\sigma_{\beta_\varphi}^2$) | 0.001 | 0.001 | 0.001 | [0.001; 0.002] |
| Variance in individual deviations from $\bar{\beta}_{\Delta\varphi}$ ($\sigma_{\beta_{\Delta\varphi}}^2$) | 0.138 | 0.139 | 0.137 | [0.099; 0.175] |
| Variance in annual deviation from $\bar{\mu}$ ($\sigma_{\mu,y}^2$) | 0.174 | 0.160 | 0.166 | [0.101; 0.292] |

Note: When the 95%CRI are above or below 0, the probability that the focal effect is respectively positive or negative is >97.5%. Values are on the logit scale. Graphical representations of the posterior distributions are in Supplemental material H.

Meanwhile, the mean effect of the defined temperature anomaly covariate (β^-^*_T_*) was negative, implying that nest completion probabilities were higher in seasonal periods when the preceding 20 days were colder than their long-term average, and hence that part of the among-year variation in phenology can arise from among-year variation in temperatures. Here, there was no detectable among-individual variation: the posterior mode of *σ*^2^ was effectively on zero (Table 1; Supplemental material H).

### Derived inferences

Transformation of the predicted liability responses (resulting from the estimated LRN parameters) into daily nest completion probabilities, ψ^-^*_d_*_,*y*_, and corresponding distributions of completion dates and proportions of non-breeders *Ω_y_*, reveals some notable conceptual and empirical insights, even given our illustrative environmentally-simplistic HM-PTM. First, the ψ^-^*_d_*_,*y*_ values are all small. On average across all years, ψ^-^*_d_*_,*y*_ peaks at only 0.081 [0.020; 0.167] (mean [95%CRI]) on day 139 [136; 144] (Figure 4A). Hence, there are no strongly predicted specific dates on which populations (or individuals) achieve the focal event (i.e. nest completion), but rather small successive daily probabilities of state switches. Correspondingly, the cumulative completion probabilities imply that the resulting peak nest completion date occurs considerably earlier than the peak nest completion probability (on day 129 [129, 129], figure 4B). This difference arises because, as the season progresses, there are fewer remaining individuals that have not yet achieved nest completion. The frequencies of new individuals that switch state therefore decrease, even if ψ^-^*_d_*_,*y*_ increases. The (HM-)PTM hence reveals how, and the degree to which, most individuals express the state switch before the date on which their switching probability would be highest.

**Figure 4.**
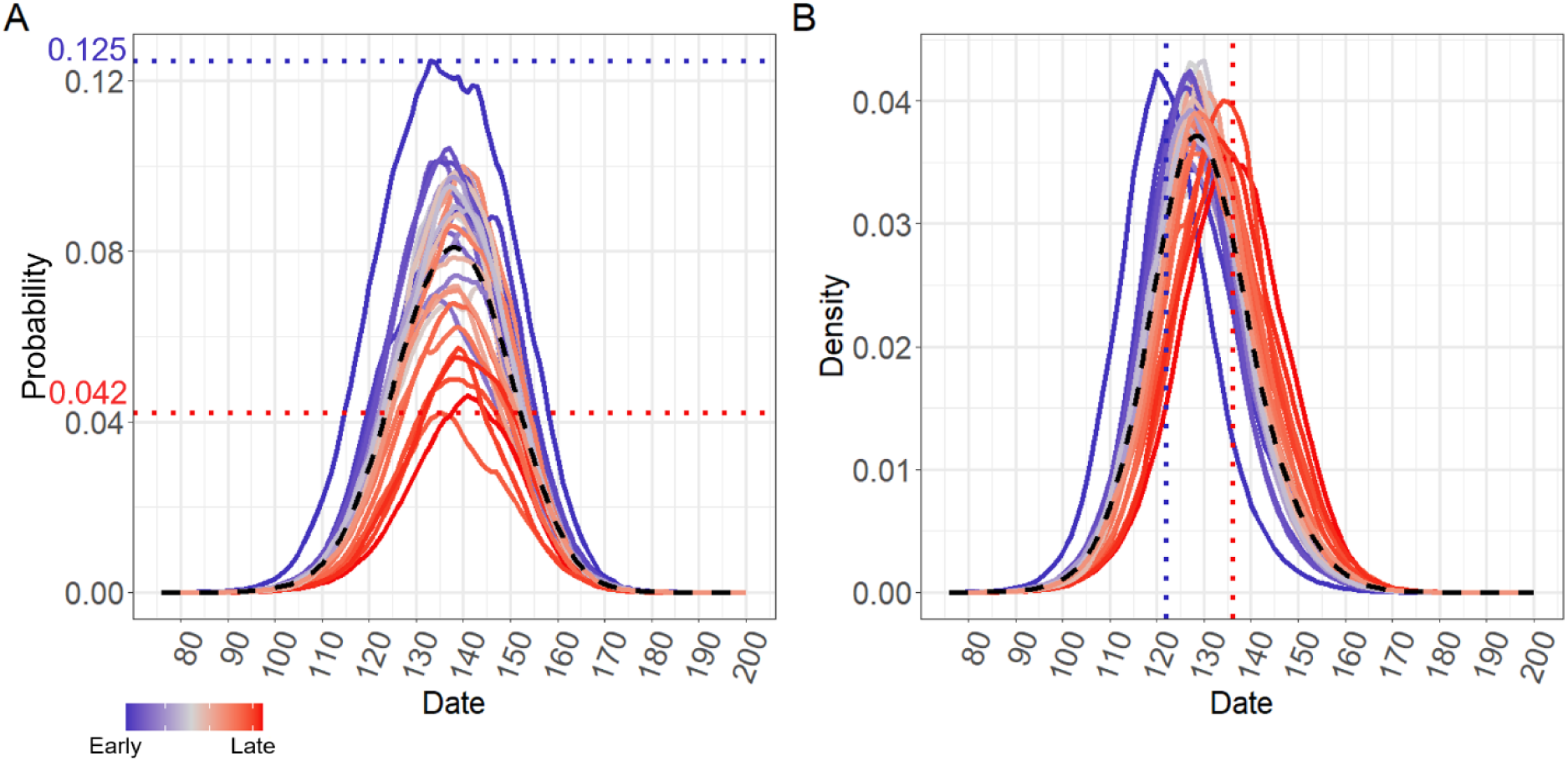
Annual daily trajectories of (A) nest completion probability, and (B) resulting annual nest completion date distributions. Black dashed lines show the grand means across all years. Coloured lines represent the posterior means for each study year (given its daily temperature anomalies*T_d_*_,*y*_ and annual deviation from the intercept *δ_μ_*_,*y*_). For visual clarity, 95%CRI are not shown. The colour gradient indicates early (blue) vs late (red) years defined from the model predictions. The dotted blue and red lines indicate the highest and lowest (A) probabilities of nest completion and (B) peaks of nest completion dates (122 and 136 respectively).

Second, the seasonal trajectory of ψ^-^*_d_*_,*y*_, and the resulting distribution of nest completion dates, varied substantially among years, stemming from variation in *T_d_*_,*y*_ and among-year variance in the LRN intercept *μ* (Equation 5, Table 1). Across the 32 study years, the lowest and highest peak values of ψ^-^*_d_*_,*y*_ were 0.042 [0.000; 0.056] and 0.125 [0.071; 0.500], with higher peak values typically associated with steeper increases and decreases in ψ^-^*_d_*_,*y*_ (Figure 4A). These patterns clearly matched the gradient from early to late phenology evident in the annual distributions of nest completion dates (colour gradient on Figure 4). Corresponding peaks of annual nest completion dates varied between ordinal date 122 [120; 123] and 136 [134; 137] (i.e. a two-week spread, Figure 4B). Moreover, the date distribution was typically narrower (and hence the density peak was higher) in years with earlier rather than later nest completion (95% of nest completion dates fell within ca. 41 and 45 days for the earliest and latest year, respectively). Hence, these patterns show how the faster liability responses in years with earlier nest completion also resulted in greater synchrony in completion.

Third, despite the simplicity of our current illustrative model in terms of annually varying environmental covariates, comparisons of the annual distributions of nest completion dates predicted from the HM-PTM versus extracted from the raw data showed good congruence in the broad pattern of among-year variation (captured primarily by *σ*^2^, Figure 5A and B). However, the predicted mean dates are typically earlier than the observed means, by 5 days on average (range 13 days earlier to 2 days later). In general, such discrepancies would be interpreted to imply some lack of model fit. However, in the present case, they can largely be explained by the incomplete field observations. Nest completion dates inferred directly from the raw field data will be biased late, and the difference between the mean model-estimated and raw observed completion dates closely matches the known average data gap stemming from the fieldwork regime (including increased discrepancies after 2004, see Methods; Supplemental material F and J). By embedding the Hidden Markov process, the HM-PTM likely estimated the distribution of event dates better than can be directly inferred from the raw (incomplete) data. For the kittiwake dataset, this implies that, contrary to initial impression from the raw data, population-wide nest completion dates have been relatively stable across the study period, with no clearly evident directional change (Figure 5A).

**Figure 5.**
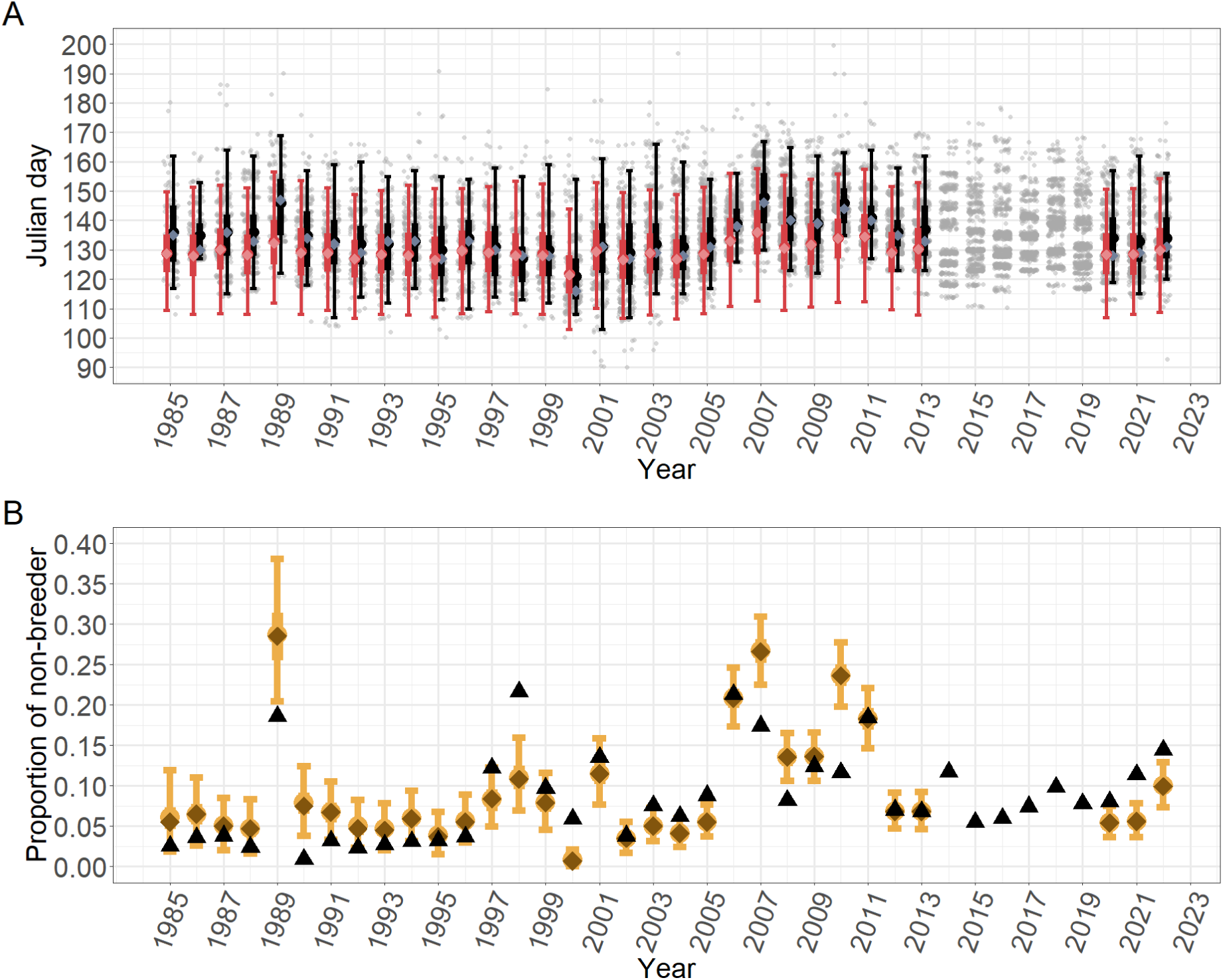
(A) Raw observations and summarised predictions of the annual distribution of breeding dates and (B) associated proportions of non-breeders in the kittiwake population. Panel A shows the mean (black points), median (grey diamonds), 50% and 95% inter-quantile ranges (respectively black thick and thin bars) of the annual observed date distributions from the raw data, and the posterior mean of the mean (pink dots), median (pink diamonds), 50% and 95%CRI (pink thick and thin bars) of the predicted annual date distributions from the HM-PTM. Each light grey points denote the first day each individual was recorded with a complete nest in each year (points are horizontally jittered for visualisation). Panel B shows the observed annual proportions of non-breeders from the raw data (black triangles) and the predicted annual mean (yellow points) and median (brown diamonds) proportions of non-breeders from the HM-PTM with their associated 50% and 95%CRI (yellow thick and thin bars).

Fourth, the predicted annual proportion of the local adult population that did not achieve nest completion, *Ω_y_*, had an overall mean of 0.09 [0.01; 0.28], which is highly consistent with the overall proportion of non-breeders directly derived from the raw data (0.09). Further, each annual *Ω_y_* varied between 0.01 [0.00; 0.02] and 0.29 [0.20; 0.38], also broadly capturing the pattern of observed among-year variation (Figure 5B). While there are some discrepancies, these fall approximately equally in both directions and hence appear unbiased. Moreover, years with low and high *Ω_y_* were also those with early and late mean nest completion dates (most notably 2000 and 2007; Figure 5). Our results therefore illustrate how the same underlying processes can generate and rationalise both phenotypic expression and phenology.

## Discussion

Predicting phenological responses to changing seasonality requires models that adequately capture realistic forms of individual responses to seasonal environmental variation and change (Hernández-Carrasco et al., 2025; Keogan et al., 2018; Wolkovich et al., 2012). Yet, by focusing on outcomes rather than temporally dynamic processes, invoking fixed ‘cue’ windows, and accepting non-random exclusion of individuals, the common approach of directly analysing variation in observed event dates could generate incomplete and misleading patterns of variation, undermining predictions (Chmura et al., 2019; Gienapp et al., 2005; Inouye et al., 2019; Kharouba and Wolkovich, 2020; Reid and Acker, 2022a). These impediments could contribute to widely documented discrepancies between expected and observed plastic and micro-evolutionary responses (Biquet et al., 2022; Bonamour et al., 2017; Chmura et al., 2019; Keogan et al., 2018; Kharouba and Wolkovich, 2020; Merilä et al., 2001; Pujol et al., 2018). To circumvent these challenges, our conceptualisation, implementation and application of a broadly-applicable Phenological Threshold Model (PTM) illustrates how explicitly formulating phenology as a seasonally dynamic state-switch can yield multiple new insights. These include magnitudes of among-individual variation underlying plastic responses to variable and invariant components of seasonal progression and resulting probabilistic distributions of phenological events, which in turn can jointly rationalise annual distributions of event dates and frequencies of non-expression.

### General principles and advances

The PTM constitutes an application of threshold trait theory to phenology, where observed event dates correspond to the moment when underlying individual and environmental characteristics combine to generate switches between alternative life-history states (Reid and Acker, 2022a,b). Analogous frameworks are established for applications in medicine (Falconer, 1965), developmental phenology (e.g., growing degree days in plant development models; Chuine and Régnière, 2017; Moore and Remais, 2014), and survival (e.g., time-to-death models; (Tutz and Schmid, 2016). We demonstrate how these concepts can be usefully employed to quantify among-individual variation in life-history plasticity, specifically in the key area of changing phenology in wild populations (Donohue et al., 2015; Gienapp et al., 2005; Reid and Acker, 2022a).

Here, there is a highly productive tradition of directly analysing dates as continuously distributed phenotypic traits (Bonamour et al., 2019; Germain et al., 2016; Inouye et al., 2019; Nussey et al., 2007; Reed et al., 2009). This approach is highly appropriate for some core purposes, for example for simply testing whether mean event dates (e.g. flowering, breeding or migration dates) differ between age-classes, habitats or populations. However, to address subsequent questions of why any differences arise, the PTM could provide new insights and alleviate key limitations of traditional date-based analyses. Specifically, by focusing on latent population and individual responses to longitudinal seasonal environmental progression, the PTM facilitates formulation and test of mechanistic hypotheses concerning biological responses to seasonally dynamic environmental drivers which do and/or do not vary among years. Key outputs comprise parameters that describe among-individual variation in intercepts and/or slopes of the underlying multidimensional latent reaction norm (LRN), and resulting individual-level and population-level probabilities of switching state within or throughout any focal period of seasonal progression. This can be achieved without defining a fixed annual ‘cue’ window that should precede all observed events, while retaining individuals that did not express the focal event and hence do not have any observed date (e.g., non-breeders, non-migrants) in any or all focal years. Further, we demonstrate how embedding a Hidden Markov model within the PTM can remove biases stemming from intermittent rather than complete observations of individuals’ seasonal state progression, potentially improving estimates of realized phenology. Indeed, the HM-PTM allows unbiased parameter estimates to be retrospectively obtained from existing data on observed event dates, even given incomplete field observations (Supplemental material J).

To illustrate these concepts, our application to annual kittiwake nest completion dates shows how even a (deliberately) simple illustrative HM-PTM with minimal environmental covariates can yield thought-provoking insights concerning the degree to which populations could harbour potential to rapidly respond to changing seasonal environments, or else exhibit key constraints. We estimated substantial liability-scale responses to fundamental aspects of seasonal progression, comprising daylength (*φ*) and the daily difference in daylength (Δ*φ*). Here, population mean responses concur with detailed physiological studies on hormonal cycles regulated by photoperiod, which have shown that long daylength induces gonadal growth and behaviours associated with breeding activities (e.g., territory defence, courtship, nest building), while long and decreasing photoperiod progressively causes gonadal regression and terminates the breeding season (Ball and Balthazart, 2003; Dawson, 2015; Dawson et al., 2001; Watanabe et al., 2007). Yet, importantly, we also estimated non-zero among-individual variance in sensitivity to *φ* and Δ*φ*. Such variation is required to achieve newly adaptive plastic and/or micro-evolutionary shifts in responses to photoperiod, and resulting phenological shifts, in the context of changing seasonality (Dawson, 2015; Hahn et al., 2025). This potential cannot be quantified by standard analyses of event dates, which rely on covariation between dates and among-year environmental variation within fixed ‘cue’ windows. This approach is nullified for daylength, because the seasonal progressions of *φ* and Δ*φ* do not vary among years.

Yet, the estimated among-individual variance in sensitivity to daylength, σ^2^_μϕ_, while non-zero, is of a magnitude that would generate little variation in switch date. Our conceptual construction (Figure 1) highlights why this might necessarily be the case, at least for populations with narrow annual windows for breeding or other events (e.g., due to narrow peaks of environmental suitability; Jacobs and Wingfield, 2000; Shultz et al., 2009; Whelan et al., 2022). Here, substantial among-individual variance in slope would generate substantial (and unrealistic) spread of event dates within each year. Since very early or late event dates might commonly imply low fitness, slopes (and associated intercepts) that generate such outcomes would be strongly selected against, and hence could be rapidly eliminated. This constraint could potentially contribute to explaining why micro-evolution of photoperiod-driven phenological change may be very slow (Hahn et al., 2009; Jacobs and Wingfield, 2000; Wingfield, 2008).

Our HM-PTM also estimated non-zero liability-scale responses to temperature anomalies, as quantified over a 20-day rolling window. This concurs with physiological studies indicating that birds perceive and integrate seasonal progression through photoperiod, then fine-tune their phenology in line with stochastic environmental variables (Ball and Balthazart, 2003; Dawson et al., 2001). While these points were previously widely suggested in phenological studies of wild populations (e.g. Charmantier et al., 2008; Chauveau et al., 2025; Keogan et al., 2022; Visser et al., 2009), links between stochastic seasonal environmental progressions and event dates, while controlling for photoperiod, have not previously been quantified. This is because, unlike the PTM, standard date analyses do not quantify effects of within-season environmental variation and change (Chmura et al., 2019; Frederiksen et al., 2004). This approach consequently precludes full clarity on how seasonally dynamic environmental variables effectively drive seasonal trajectories of individual liabilities that underpin phenology, preventing identification of constraints and limiting future predictions (Chmura et al., 2019).

Our current objective was to provide illustrative rather than comprehensive investigation of relationships between kittiwake nest completion and seasonal environmental variation. Our defined temperature anomaly and daylength variables could be correlated with other variables that might have more direct causal effects (e.g., rainfall, spring phytoplanktonic bloom, food availability; (Cullen, 1957; Shultz et al., 2009; Suryan et al., 2002). Clearly, future studies designed to test hypotheses explaining (lack of) phenological changes in any system will require longitudinal (within-season) data on appropriate suites of ecologically-relevant environmental drivers that are postulated to affect the LRN intercept and/or slopes (e.g., air, water or sea temperatures, daily insolation, ice break-up, or bursts of lower trophic level prey, Calle et al., 2010; Frederiksen et al., 2007; Tessmar-Raible et al., 2011; Visser et al., 2009; Whelan et al., 2022). Such work could exceed the performance of our current analyses in capturing observed among-year variation in phenology through explicit predictors rather than predominantly through random year effects on μ, thereby facilitating prediction of future outcomes.

### Perspectives and opportunities

Having demonstrated the (HM)-PTM concept, implementation and basic application, multiple new biological insights can now be achieved through application to diverse systems, requiring careful analytical decisions and further conceptual and technical developments. The PTM can be fitted to any type of longitudinal dichotomous data involving any time grain, including natural or experimental systems where focal events occur on timeframes of seconds, minutes or hours rather than days, weeks or months. The HM component could handle observation failures occurring on any such timeframe, including experimental studies where observations occur at set intervals (and hence effectively record whether state-switches have occurred since the previous observation). Further evaluations of (HM-)PTM performance and parameter identifiability under different circumstances will then be required, including when numbers of observed switches per individual are inherently limited (e.g. in short-lived or elusive organisms).

Alongside careful specification of appropriate environmental covariates, analyses must also specify the timeframes over which such variables are postulated to act. Impacts could be effectively instantaneous (e.g. where daily environments directly affect daily liabilities), or could be time-lagged, or averaged, or accumulated over some specified time period (as in our rolling mean formulation, or commonly captured as degree days of warming; Cayton et al., 2015; Chuine and Régnière, 2017; Gienapp et al., 2013). The full analysed time period D must also be carefully defined, since the selected start time will affect the interpretation of variance in the liability scale intercept, and the end time will affect the interpretation of individuals that did not (yet) express the focal event. While it enables explicitly modelling the dynamics of seasonal progression, the (HM-)PTM formulation does not preclude modelling effects of preceding environmental ‘cues’, if desired. This could be achieved by modelling effects of environmental variables measured at fixed preceding timepoints on the LRN intercept. Hence, overall, (HM-)PTMs could be directly formulated based on existing knowledge of system ecology and/or physiology, or be deployed to statistically test hypotheses on influential covariates and their pace of effect. For example, appropriate D could be defined from historical records of early or late events or oestrus/anoestrus cycles (Dawson et al., 2001), or from hypotheses on key resources and environmental conditions experienced before versus during breeding seasons could be tested (e.g., in capital versus income breeders;(Stephens et al., 2014).

More specific mechanistic hypotheses could also be tested, including potential trade-offs, non-linearities and interactions that could facilitate or constrain rapid phenological changes. For example, strong negative intercept-slope covariances, or non-linear LRNs, could further shape the translation of seasonal progression into the phenology of life-history events (Reid and Acker, 2022a). Interactions between variable and invariant drivers of seasonal progression could be explicitly quantified, for example photoperiod by temperature interactions such as regulate budburst in birch, *Betula pubescens* (Caffarra et al., 2011), or photoperiod by food interactions such as regulate breeding in red crossbill, *Loxia curvirostra* (Hahn, 1998). New insights into the eco-physiological basis of structured life-history variation could then follow. For example, (HM)-PTMs could include variables such as individual age, life-history stage, breeding experience, social context and/or sub-population in addition to (or in interaction with) environmental variables alongside any other desired fixed or random effects. Such analyses could reveal, for example, whether widely observed effects of age, breeding experience, mate retention and/or sub-population on reproductive phenology (Auld and Charmantier, 2011; Caro et al., 2009; Coulson and White, 1956, 1959; Lambrechts et al., 1997; Lewis et al., 2025) stem from differing sensitivities to seasonal progression (manifesting as differing LRN slopes) and/or differing starting points (manifesting as differing LRN intercepts), versus simply from differing environmental variables.

Modelling such effects should further improve joint estimation of resulting date distributions and of non-event (e.g. non-breeding) frequencies. Such models could also explain how skewed annual date distributions, which are commonly observed in nature (Bonamour et al., 2017; Moyes et al., 2011), may emerge from non-linear seasonal trajectories of underlying liability responses (Reid and Acker, 2022a). Finally, the (HM-)PTM approach could underpin new analyses of phenological mismatches and potential evolutionary constraints on phenology. Quantifying relationships linking underlying genetic variation to event dates and how they vary with the environment could facilitate testing hypotheses on how selection operating through dates given phenological mismatches ultimately acts on LRN intercepts and slopes and hence results in evolutionary change or stasis. More robust predictions of phenological responses to changing seasonality could thereby be achieved, improving understanding of population resilience in the face of climate change.

## Data and Code Availability

Data and codes to reproduce all simulations, analyses and figures are provided in a publicly available external repository (Anonymous, 2026). Private access link for reviewing: http://datadryad.org/share/LINK_NOT_FOR_PUBLICATION/NBkdo7jIj_zxEkMC9-DeI-Ewi3Cypno0RNnUUIM8xvE.

## Ethics statement

All fieldwork was licensed by the Centre de Recherches sur la Biologie des populations d’Oiseaux (‘CRBPO’, Museum National d’Histoire Naturelle, Paris, France), and carried out in accordance with standard animal care protocols approved by the CRBPO.

## Author contributions

PA and JMR conceived the overarching objectives. PA devised the quantitative framework and coded the statistical models. VG and PA coded and analysed the simulations, with input from JMR. VG adapted the models for analyses of the kittiwake data and conducted the analyses, with input from PA, JMR, and EC. EC collected the data. VG drafted the manuscript, with assistance from JMR and PA. All authors contributed to manuscript review and editing.

## Acknowledgements

This work was supported by ISblue project, Interdisciplinary graduate school for the blue planet (ANR-17-EURE-0015) and co-funded by a grant from the French government under the program ‘Investissements d’Avenir’ embedded in France 2030, the SEA-Life programme of the French National Centre for Scientific Research (‘CNRS INEE’, Ecology and Evolution Institute), the marine observatory of the European Institute for Marine Studies (‘IUEM’, UAR 3113), and the Langlois Foundation. JMR and PA were partially supported by the European Research Council (Advanced Grant EcoEvoMove 101140637 awarded to JMR). Views and opinions expressed are however those of the authors only and do not necessarily reflect those of the European Union or the European Research Council Executive Agency. Neither the European Union nor the granting authority can be held responsible for them.

