## Supplemental Material A to J for "Conceptualising and quantifying phenological plasticity through latent responses to seasonal environmental progression"

This supplemental material provides further details on the study system, datasets and results, along with mathematical formulas underpinning our models. The supplemental material contains A-J sections, Supplemental Figures B, D, E, F1-3, G1-2, H, I and J, and Supplemental Tables A, E, H1-2.

### Supplemental material A: Table with symbols and definitions

Table A summarises all key symbols and associated definitions from the main text.

*Table A. List of symbols and associated definitions.*

| Symbol | Definition |
|---|---|
| $\boldsymbol{i}$ | An individual within the study population. |
| $\boldsymbol{y}$ | A focal year among the monitored years, during which individual *i* might switch state and hence express the focal phenotype. |
| $\boldsymbol{d}$ | One timestep (i.e., day) through the focal season within year *y*. |
| **D** | Duration of the annual study season (in days). |
| $\boldsymbol{\ell_{i,d,y}}$ | Liability of individual *i* on day *d* in year *y*. This is the latent continuous variable that summarizes the joint biological effect of all genetic, individual, and environmental factors underlying the expression of the dichotomous phenotypic states (A or B) which define the focal phenological event. |
| **θ** | Threshold, the liability value above which an individual's seasonal state switch (from state A to B) occurs. |
| $\boldsymbol{x_{d,y}}$ | Environmental covariate that may change among days *d* within any year *y*. This pattern of change can vary (e.g., temperature), or not (e.g., photoperiod), among years. Conversely, $x_{d,y}$ could also remain constant across days but vary among years (e.g. density in a closed population). |
| **LRN** | Latent Reaction Norm, the function linking the liability (the response $\ell_{i,d,y}$) to one or multiple environmental covariates $x_{d,y}$ (the driving variables). |
| $\boldsymbol{\bar{\mu}}$ | Population mean intercept of the LRN. |
| $\boldsymbol{\bar{\beta}_x}$ | Population mean slope of the LRN, representing mean sensitivity of the population to the focal environmental covariate $x$. |
| $\boldsymbol{\delta_{\mu,i}}$ | Individual deviation from $\bar{\mu}$, where $\delta_{\mu,i} \sim \mathcal{N}\left(0, \sigma_{\mu}^2\right)$, implying among-individual variation in $\mu$. |
| $\boldsymbol{\delta_{\beta_x,i}}$ | Individual deviation from $\bar{\beta}_x$, where $\delta_{\beta_x,i} \sim \mathcal{N}\left(0, \sigma_{\beta_x}^2\right)$, implying among-individual variation in sensitivity to $x$. |
| $\boldsymbol{s_{i,d,y}}$ | Phenotypic state (either A or B) of individual *i* on day *d* in year *y*. Note that the switch from A to B is presently assumed to be irreversible. |

| | |
|---|---|
| $\boldsymbol{H_{i,y}}$ | Observed phenological sequence of phenotypic states on each successive day from 1 to D, for individual *i* in year *y*. |
| $\boldsymbol{a_{i,y}}$ | Last day individual *i* was observed in state A in year *y*. |
| $\boldsymbol{b_{i,y}}$ | First day individual *i* was observed in state B in year *y*. This value may not exist for some individuals in some years. |
| $\boldsymbol{\chi_{i,t,y}}$ | Probability (under the model) that individual *i* switches state in year *y* on any day within D, from after day *t* to $b_{i,y}$, conditional on the switch having not yet occurred on $t$. Note that $\chi_{i,t,y}$ is defined strictly by the model parameters given any hypothetical *t* and $b_{i,y}$. |
| $\boldsymbol{\Psi_{i,d,y}}$ | Probability that individual *i* expresses the focal life-history event (i.e. switches from phenotypic state A to B) on day *d* in year *y*, conditional on not having switched yet in the focal season (i.e. prior to *d* in *y*). Applying the link function to $\Psi_{i,d,y}$ gives the underlying expected liability on the link scale (see **Supplemental material C**). |
| $\boldsymbol{\bar{\Psi}_{d,y}}$ | Mean population-wide probability that individuals will express the focal life-history event (i.e. switch phenotypic state) on day *d* in year *y*, among individuals that have not previously switched during the focal year. This is derived as a posterior prediction from the fitted PTM. |
| $\boldsymbol{\Omega_y}$ | Proportion of individuals that do not switch state in year *y*. This is derived as a posterior prediction from the fitted PTM. |

29

#### Supplemental material B: Data and analysis structures

Here, we summarise decisions taken for our current analyses, and highlight general decision points for analysing discrete phenotypic traits with the threshold trait framework.

Our current analyses focus on the case of a dichotomous trait that is not reversible within years (red dotted frame in Figure B). This matches the common conceptualisation of phenological events (i.e., with only one phenological switch possible within a year). This envisages that, for example, post-breeding (including following early failure) is not the same biological state as pre-breeding. This assumption of non-reversibility is necessary for our illustrative analyses of the kittiwake data (bold frame, Figure B) because of the incomplete field observations that generate gaps within individuals' observed state sequences. Obtaining unbiased estimates of the underlying LRN parameters then requires the Hidden Markov model component, where our current algorithm assumes that individuals cannot reversibly switch between phenotypic states within or encompassing unobserved timepoints. This assumption could potentially be relaxed in future analyses designed for systems and traits that are readily reversible (e.g. behavioural states). In addition, the assumption of non-reversibility enables back-filling unknown phenotypic states with the pre-switch state before the switch happens. For example, in a dichotomous trait with successive potential phenotypic states A and B, NAs between successive As within a state sequence can be replaced with As.

In the absence of any state sequence gaps, exploratory analyses showed that the PTM can readily handle cases where the phenological trait is reversible or non-reversible. In fact, these analyses indicated that the PTM performs even better in estimating LRN parameters for reversible traits (i.e. more accurate and precise estimates). This is probably because there are more switching events, and a lack of reverse switching soon after any one switching also informs on the reaction norm slope.

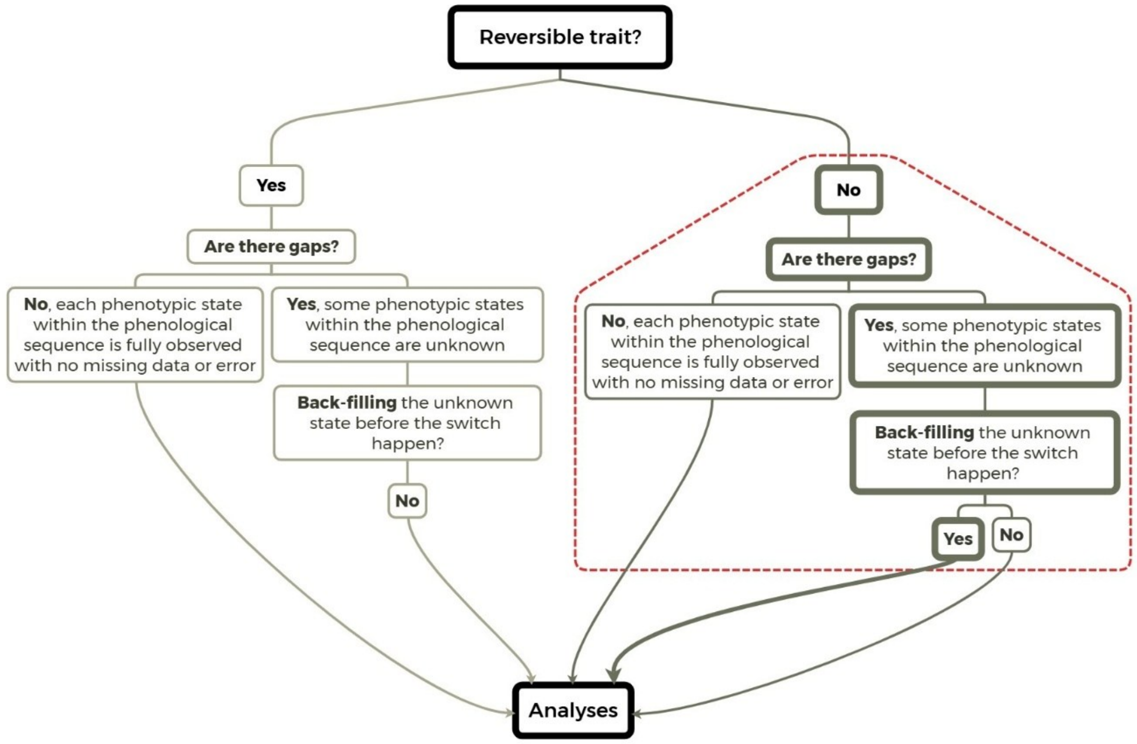

*Figure B. Decision tree of choices embedded in applications of the Phenological Threshold Model (PTM). The red dotted frame highlights the path taken for our current methodological developments and analyses. Bold rectangles correspond to the path for modelling the kittiwake breeding phenology.*

### Supplemental material C: Formulating a GLMM to estimate LRN parameters

Here, we detail the transformation linking the underlying liability reaction norm (LRN) parameters to the dichotomous phenotypic scale, allowing estimation of the LRN parameters from observed binary state sequences.

Under the PTM framework, the focal state transition of an individual *i* in year *y* occurs when its liability on day $d$, $\ell_{i,d,y}$, exceeds a threshold, θ. Following our reaction norm framework (main Figure 1), we express $\ell_{i,d,y}$ as a linear function encompassing a general intercept plus the effects of environmental covariates, including among-individual variation in these parameters and a within-individual daily residual. In the case where there is only one environmental covariate $x_{d,y}$, $\ell$ is thus formulated as:

$$\ell_{i,d,y} = \bar{\mu}^* + \delta^*_{\mu,i} + (\bar{\beta}^*_x + \delta^*_{\beta_x,i}).x_{d,y} + \varepsilon^*_{i,d,y}, \qquad \text{(C1)}$$

where * denotes parameter values on the original latent liability scale; $\bar{\mu}^*$ is the population mean liability intercept; $\bar{\beta}^*_x$ is the population mean regression slope on an environmental variable $x$; $\delta^*_{\mu,i}$ and $\delta^*_{\beta_x,i}$ are the individual deviations from $\bar{\mu}^*$ and $\bar{\beta}^*_x$, respectively, and $\delta^*_{\mu,i} \sim \mathcal{N}\left(0, \sigma^{*2}_{\mu}\right)$ and $\delta^*_{\beta_x,i} \sim \mathcal{N}\left(0, \sigma^{*2}_{\beta_x}\right)$ where $\sigma^{*2}_{\mu}$ and $\sigma^*_{\beta_x}$ are their respective variances; and $\varepsilon^*_{i,\,d,y}$ is the residual, which can be assumed to follow a logistic distribution with location 0 and scale $s^*$, $\mathcal{L}(0, s^*)$.

Ultimately, we want to formulate the likelihood for observations of the dichotomous life-history state, and thus formulate the probability that individual *i* expresses state B on day *d* in year *y* (conditional on not having switched yet), $\Psi_{i,d,y}$, as a function of the liability-scale parameters. Given the threshold trait model (see explanations in main text), $\Psi_{i,d,y}$ is the probability that $\ell_{i,d,y}$ exceeds θ:

$$\Psi_{i,d,y} = P\big(\ell_{i,d,y} > \theta\big). \tag{C2}$$

Moreover, letting $\eta^*_{i,d,y} = \bar{\mu}^* + \delta^*_{\mu,i} + (\bar{\beta}^*_x + \delta^*_{\beta_x,i}).x_{d,y}$ (i.e. $\eta_{i,d,y}$ is the linear predictor of $\ell_{i,d,y}$), we then have:

$$\ell_{i,d,y} \sim \mathcal{L}(\eta^*_{i,d,y}, s^*). \tag{C3}$$

However, $\ell$ is a latent variable of unknown unit that cannot be measured or observed. Therefore, θ is unknown, and the link to phenotype inference on $\Psi_{i,d,y}$ cannot be made directly through Equation C2. To circumvent this problem, we employ the following standardization, whereby liabilities are threshold-centered and scaled by $s^*$:

$$\ell'_{i,d,y} = \frac{\ell_{i,d,y} - \theta}{s^*}. \tag{C4}$$

The standardized liability values $\ell'_{i,d,y}$ are therefore defined as distances to the threshold and their unit is the scale of the liability residuals. This yields a reparameterization of our initial linear model on the original liability scale (Equation C1), to a standardized scale where the threshold is 0, and the residuals have a scale of 1 (and hence follow a standard logistic distribution; Acker & Reid, 2025). We thus now have:

$$\Psi_{i,d,y} = P\big(\ell'_{idy} > 0\big). \tag{C5}$$

$$\ell'_{idy} \sim \mathcal{L}(\eta'_{i,d,y}, 1). \tag{C6}$$

$$\eta'_{i,d,y} = \frac{\eta^*_{i,d,y} - \theta}{s^*}. \tag{C7}$$

Accordingly, $\Psi_{i,d,y}$ can be expressed using the cumulative distribution function of the standard logistic distribution (which is the inverse of the logit function) as:

$$\Psi_{i,d,y} = \mathrm{logit}^{-1}(\eta'_{i,d,y}). \tag{C8}$$

We thereby retrieve the regression Equation of a binary GLMM with a logit link as formulated in Equation 1 of the main text:

$$logit\left(\Psi_{i,d,y}\right) = \eta'_{i,d,y} = \bar{\mu} + \delta_{\mu,i} + \left(\bar{\beta}_x + \delta_{\beta_x,i}\right).x_{d,y}. \quad \text{(C9)}$$

where: $\bar{\mu} = \frac{\bar{\mu}^* - \theta}{s^*}$, $\bar{\beta}_x = \frac{\bar{\beta}_x^*}{s^*}$, $\delta_{\mu,i} = \frac{\delta^*_{\mu,i}}{s^*}$, $\delta_{\beta_x,i} = \frac{\delta^*_{\beta_x,i}}{s^*}$, $\sigma_\mu^2 = \frac{\sigma_\mu^{*2}}{s^{*2}}$ and $\sigma_{\beta_x}^2 = \frac{\sigma_{\beta_x}^{*2}}{s^{*2}}$ (Acker and Reid, 2025). Hence, by fitting a binary GLMM to the observed dichotomous state data, we can estimate all LRN parameters on a standardized liability scale (the logit scale). As noted above, on this scale, the latent liability residuals follow a standard logistic distribution, and thus the residual variance is $\frac{\pi^2}{3}$. Hence the unit is $\frac{\sqrt{3}}{\pi}$ times the standard deviation of the liability residuals (which cannot be estimated and serves as the benchmark to measure variation in liability).

This PTM, without any embedded Hidden Markov component, can be readily fitted in standard statistical softwares in either Bayesian or frequentist frameworks.

### Supplemental material D: Backward recursion

Here, we detail the backward recursion applied to handle missing data and eliminate bias that would otherwise result from incomplete field observations.

The backward recursion is a simple algorithm that facilitates calculation of the probability $\chi_{i,a_{i,y},y}$ that individual *i* switched to state B on any particular day after the last day it was observed in state A, $a_{i,y}$, if it was ever observed in state B (i.e. if $b_{i,y}$ exists). More precisely, $\chi_{i,a_{i,y},y}$ is the sum of the probabilities of all possible underlying state sequences from $a_{i,y}$ to $b_{i,y}$, through the days when the individual's state was not observed (Figure D). The recursion uses the simple property that, for any day $t$ from $a_{i,y}$, the probability that *i* switches on any day after $t$ (i.e. $\chi_{i,t,y}$) is simply the probability that *i* switches on *t*+1 (conditional on not having switched yet, i.e. $\Psi_{i,t+1,y}$) plus the probability that it did not switch on *t*+1 but switched on any one day after *t*+1 (i.e. $(1-\Psi_{i,t+1,y})\chi_{i,t+1,y}$; Equation 4 of the main text, Figure D). Since we know that $\chi_{i,t,y}$ = 0 if $t \geq b_{i,y}$ (because then *i* has already switched, and the focal life-history trait is irreversible within a given year), we can start from $t = b_{i,y}$ and proceed by sequentially calculating $\chi_{i,t,y}$ for each preceding day in reverse chronological order down to $t = a_{i,y}$.

For example, with a gap of 4 days (i.e. Figure D), we start with $t = a_{i,y} + 5 = b_{i,y}$, with $\chi_{i,b_{i,y},y} = 0$. Then, for each day until $t = a_{i,y}$ (or $t = b_{i,y} - 5$):

$$\begin{aligned}\chi_{i,b_{i,y}-1,y} &= \chi_{i,a_{i,y}+4,y} \\ &= \Psi_{i,b_{i,y},y} + \left(1-\Psi_{i,b_{i,y},y}\right).\chi_{i,b_{i,y},y} \qquad \text{(D1)} \\ &= \Psi_{i,b_{i,y},y}\end{aligned}$$

$$\chi_{i,b_{i,y}-2,y} = \chi_{i,a_{i,y}+3,y} \qquad \text{(D2)}$$

$$= \Psi_{i,b_{i,y}-1,y} + \left( 1 - \Psi_{i,b_{i,y}-1,y} \right) . \chi_{i,b_{i,y}-1,y}$$

$$= \Psi_{i,b_{i,y}-1,y} + \left( 1 - \Psi_{i,b_{i,y}-1,y} \right) . \Psi_{i,b_{i,y},y}$$

$$\chi_{i,b_{i,y}-3,y} = \chi_{i,a_{i,y}+2,y}$$

$$= \Psi_{i,b_{i,y}-2,y} + \left( 1 - \Psi_{i,b_{i,y}-2,y} \right) . \chi_{i,b_{i,y}-2,y} \qquad \text{(D3)}$$

$$= \Psi_{i,b_{i,y}-2,y} + \left( 1 - \Psi_{i,b_{i,y}-2,y} \right) . \left( \Psi_{i,b_{i,y}-1y} + \left( 1 - \Psi_{i,b_{i,y}-1,y} \right) . \Psi_{i,b_{i,y},y} \right)$$

$$\chi_{i,b_{i,y}-4,y} = \chi_{i,a_{i,y}+1,y}$$

$$= \Psi_{i,b_{i,y}-3,y} + \left( 1 - \Psi_{i,b_{i,y}-3,y} \right) . \chi_{i,b_{i,y}-3,y}$$

$$\text{(D4)}$$

$$= \Psi_{i,b_{i,y}-3,y} + \left( 1 - \Psi_{i,b_{i,y}-3,y} \right) . (\Psi_{i,b_{i,y}-2,y} + \left( 1 - \Psi_{i,b_{i,y}-2,y} \right) . \left( \Psi_{i,b_{i,y}-1y} + \right.$$

$$\left. \left( 1 - \Psi_{i,b_{i,y}-1,y} \right) . \Psi_{i,b_{i,y},y} \right))$$

$$\chi_{i,b_{i,y}-5,y} = \chi_{i,a_{i,y},y}$$

$$= \Psi_{i,b_{i,y}-4,y} + \left( 1 - \Psi_{i,b_{i,y}-4,y} \right) . \chi_{i,b_{i,y}-4,y}$$

$$\text{(D5)}$$

$$= \Psi_{i,b_{i,y}-4,y} + \left( 1 - \Psi_{i,b_{i,y}-4,y} \right) . (\Psi_{i,b_{i,y}-3,y} + \left( 1 - \Psi_{i,b_{i,y}-3,y} \right) . (\Psi_{i,b_{i,y}-2,y} +$$

$$\left( 1 - \Psi_{i,b_{i,y}-2,y} \right) . \left( \Psi_{i,b_{i,y}-1y} + \left( 1 - \Psi_{i,b_{i,y}-1,y} \right) . \Psi_{i,b_{i,y},y} \right)))$$

That way, $\chi_{i,a_{i,y},y}$ is formulated as a succession of $\Psi_{i,t,y}$ for each $t$. This recursive algorithm facilitates generic implementation of the (HM-)PTM, as it is readily coded through a simple backward 'for loop' statement (see archived code: Anonymous, 2026).

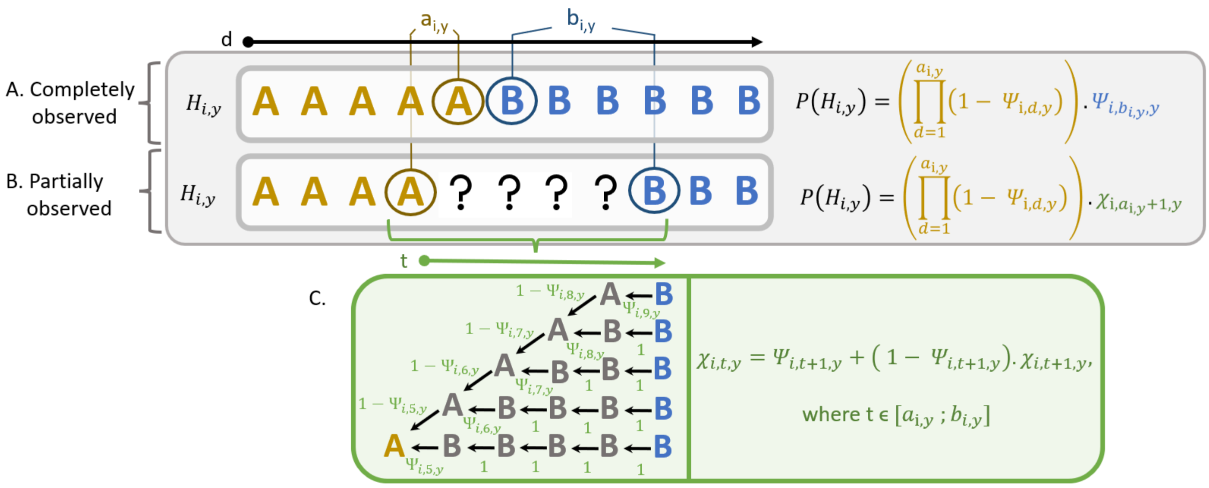

*Figure D. Phenological Threshold Model and associated state sequence data ($H_{i,y}$) when $H_{i,y}$ is (A) completely observed or (B) not completely observed, for an individual i switching state within year y. The embedded Hidden Markov model is presented in the green rectangle (C) for a non-reversible phenological trait with two alternative phenotypic states, A and B, applied for individual i, that switched state on day d within year y.*

#### Supplemental material E: Simulated datasets

We designed simulations to test the ability of the (HM-)PTM to estimate latent reaction norm parameters describing liability responses to environmental drivers of phenological events, including among-individual variances.

We first simulated two different environmental variables, $x_{d,y}$, each of which varied over 50 days (denoted *d*) within each of 6 years (denoted *y*). We thereby envisage a system where individuals' life-history sequences are repeatedly monitored over several years, facilitating estimation of among-individual variances. The environmental variables are respectively deterministically (Equation E1) or stochastically (Equation E2) linearly related to day *d* (and hence to date), where:

$$x_{d,y} = \gamma + \alpha_y . d \qquad \text{(E1)}$$

$$x_{d,y} = \gamma + \alpha_y . d + z_d \qquad \text{(E2)}$$

Here, $\gamma$ is the intercept which we set to zero; $\alpha_y$ is the slope for each year, which we drew from a uniform distribution, such as $\alpha_y \sim U(0.2\ ,\ 0.3)$; $z_d$ is an additional noise for each day *d*, which depends on the previous day's value, $z_{d-1}$. This temporally autocorrelated noise was generated using a simple auto-regressive model of order 1 (Equation E3):

$$z_d = p . z_{d+1} + \varepsilon_d \qquad \text{(E3)}$$

where the auto-regressive coefficient $p$ was set to 0.8, and the residual $\varepsilon_d$ was drawn independently for each day from a normal distribution with mean 0 and variance $0.37^2$.

We then set appropriate LRN parameter values (Table E1) which, given the simulated environmental variables, generate temporal patterns of state-switches that approximate biologically plausible variation in phenology. Specifically, we defined a negative mean intercept, $\bar{\mu}$, thereby ensuring that liability values start below the threshold at *d*=1. We defined a positive mean slope, $\bar{\beta}_x$, thereby ensuring that liabilities increase progressively as each year progresses, given that the environmental variables were defined to increase with *d*. Together, these parameters ensured that all individuals were expected to cross the threshold (and hence switch state) within the simulated 50-day period within each year. The variances of the individual deviations from $\bar{\mu}$ and $\bar{\beta}_x$ (i.e., $\sigma_\mu^2$ and $\sigma_{\beta_x}^2$ , assuming that the deviations are normally distributed with a mean of zero) were then specified to generate biologically plausible degrees of among-individual variation in the day on which the threshold was crossed, and hence the date of state-switching. Given the primary initial aim of verifying the fundamental ability of the PTM to retrieve the specified LRN parameters, including given missing observations, the specified variances only generated a small probability that individuals' liabilities would not cross the threshold.

We initially envisaged that there could potentially be a lack of identifiability, particularly between $\sigma_\mu^2$ and $\sigma_{\beta_x}^2$. We therefore constructed two sets of internally consistent parameter values with the among-individual variance in the LRN slope ($\sigma_{\beta_x}^2$) set to zero or not, which required a reparameterization of $\bar{\beta}_x$ (Table E1).

Given each of the two simulated environmental covariates (linear and non-linear) and two sets of parameters (with and without non-zero $\sigma_{\beta_x}^2$), we generated LRNs by using the specified values (Table E1) to parameterize Equation C1, drawing the individual daily residuals from a standard logistic distribution. In each of the four cases, we simulated a total of 6000 individual state sequences for each year ($H_{i,y}$), for a fully balanced dataset envisaging 1000 individuals that were all observed in each of six years (i.e. 6000 individual-years), and an unbalanced dataset in which 3706 individuals were observed in a varying number out of the six years (ranging from 1-6, also totaling 6000 individual-years). For this unbalanced dataset, the number of individuals observed per year is generated such that most individuals are resighted only once or twice (as is the case for the kittiwake data). To get the appropriate number of individuals resighted per year required to ensure a total of 6 000 $H_{i,y}$, we generated a distribution with a large sample of individuals following a Beta distribution, $Beta(0.8, 2)$, over 6 years to get the proportion of individuals resighted per year, and then spread the 6000 $H_{i,y}$ accordingly, resulting in a total of 3706 individuals. The large sample sizes of the datasets were chosen to ensure high statistical power, given our current objective of testing whether the (HM-)PTM was capable of returning unbiased parameter estimates, not of evaluating what sizes or structures of datasets would be required in practice. In each case, we used the LRNs (given the defined threshold of zero) to generate the full corresponding $H_{i,y}$ for each year. These state sequences were then either retained in full or degraded by only retaining every sixth datum (and substituting NA otherwise), thereby envisaging a fieldwork regime of observations every six days. Figure E illustrates the sequential process followed to build the total of 16 simulated datasets.

The PTM and/or HM-PTM were then fitted to the resulting simulated datasets with the individual state sequences using weakly informative priors (i.e., $\bar{\mu}$ and $\bar{\beta}_x \sim Student\text{-}t(3, 0, 2.5)$, and $\sigma_\mu$ and $\sigma_{\beta_x} \sim Half\text{-}Student\text{-}t(3, 0, 2.5)$ , with 1,000 warm-up iterations followed by 1,000 monitored iterations for each of four chains, resulting in total posterior sample sizes of 4,000.

Our current simulations were designed to provide proof of functional concept of the (HM-)PTM, also broadly envisaging the exemplar kittiwake dataset (which includes observations of many individuals in multiple years). However, our simulation procedures and codes can be used to examine (HM-)PTM performance under conditions that might be relevant for other systems and datasets, for example fewer individuals and/or few individuals that were observed in multiple years.

*Table E1. Parameter values used to generate the simulated Liability Reaction Norms (LRNs).*

| Parameter | Parameter values | |
|---|---|---|
| | **Simulations with $\sigma^2_{\beta_x} = 0$** | **Simulations with $\sigma^2_{\beta_x} \neq 0$** |
| **Mean intercept ($\bar{\mu}$)** | -13.5 | |
| **Mean slope ($\bar{\beta}_x$** | 2.2 | 4.6 |
| **Variance in individual deviations from $\bar{\mu}$ ($\sigma^2_{\mu}$)** | 6.6 | |
| **Variance in individual deviations from $\bar{\beta}_x$($\sigma^2_{\beta_x}$)** | 0 | 0.46 |

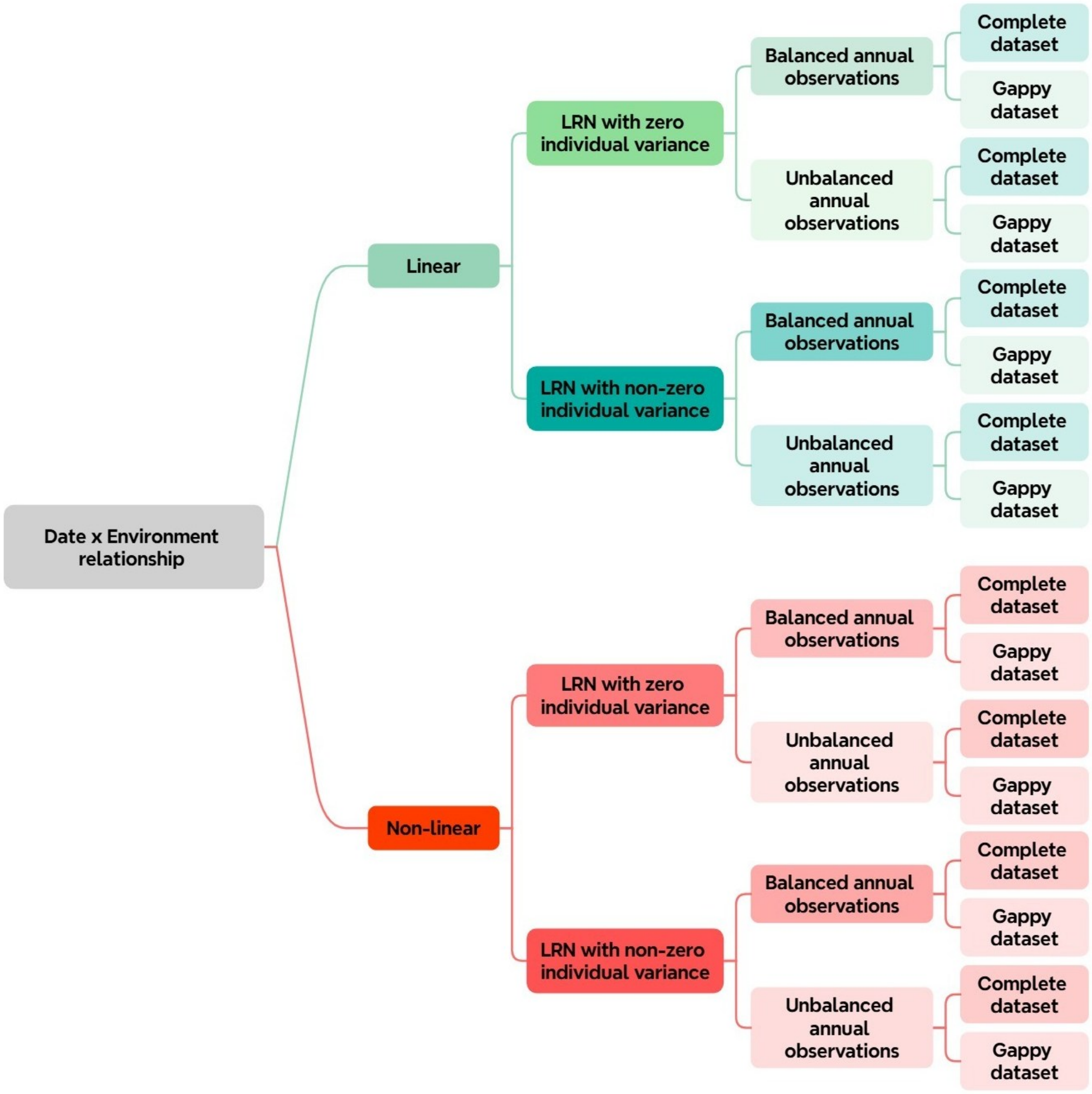

*Figure E. Flowchart illustrating the different steps to build the 16 simulated datasets.*

### Supplemental material F: Details of the kittiwake study system and collected data

The black-legged kittiwake (*Rissa tridactyla*, family Laridae) is a long-lived pelagic and migratory seabird, wintering throughout the northern hemisphere. During the breeding season (Figure F1), kittiwakes gather in colonies along North Atlantic coastlines. They form socially monogamous pairs which can last several years (Naves et al., 2005), and build nests on cliffs (Cullen, 1957; Maunder & Threlfall, 1972). A completed nest is an elaborate structure (Figure F2.A) and is considered as a reliable criterion to define physiological and behavioral maturity for reproduction. Indeed, nest completion requires both pair members to be experienced enough and coordinated to gather materials (e.g., grass, mud, sticks, seaweeds), and defend the nesting site and materials from neighboring kittiwakes (Cullen, 1957; Danchin, 1987; Maunder & Threlfall, 1972).

Since 1979, thousands of kittiwake chicks hatched in Cap Sizun (48°03'N, 4°39'W), France, have been marked with unique combinations of colored rings (Figure F2). This enables subsequent reproductive monitoring of yearly breeding activities of individuals that survived to adulthood and locally recruited. Each year, from the first bird arrivals (January-June) until their departures (in August; Figure F1), colonies are visited every few days (on average every 1 to 4 days between 1985 and 2003, and every 5 to 12 days since 2004) before the first chicks can fledge (Figure F3). Then, visits become daily until the last individual leaves the colonies. These visits allow recording ringed individuals' nest locations and phenological information (i.e., dates of observed arrival, nesting, laying, hatching, fledging and departure; Figure F1), along with breeding state (i.e., breeders, skippers; Acker et al., 2022; Cam et al., 1998).

Although monitoring started in 1979, our current analysed dataset starts in 1985. This is because the standardized protocol to collect data has been finalized in 1983, and because we only consider individuals ringed as chicks that survived to locally recruit, implying a delay (average age at first breeding is 4 years old; Danchin & Cam, 2002). Nevertheless, the available data provide a good opportunity to apply the (HM-)PTM to real biological data for numerous individuals (i.e., 3209) while covering a period (i.e., 32 years) long enough to encompass a wide range of environmental conditions.

Our current illustrative analyses focus on the trait of nest completion, which is interpretable as one measure of the time at which individuals initiate breeding and switch from non-breeding to breeding state (respectively yellow and blue in Figure F). In this kittiwake population, nest completion is a more reliable measure than observation of eggs. Indeed, nest contents cannot be observed due to limited accessibility (i.e., cliffs), and eggs are commonly laid and lost between visits due to biotic or abiotic factors (e.g., weather, nest instability, predation, accidents; Cam et al., 1998). In contrast, a completed nest is a big and longer-lasting structure that is easily seen and evaluated (Figure F2.A).

Over the study period, the number of annual observations has increased (Figure F2.A), but the visits have become more spaced (i.e. longer gaps), decreasing the precision of the phenological date recorded. Moreover, few nests (3% of the overall dataset, Figure F3.A and B) were first recorded as already completed. These occurrences were all late within the resighting season (Figure F3.C), and hence included in our analyses.

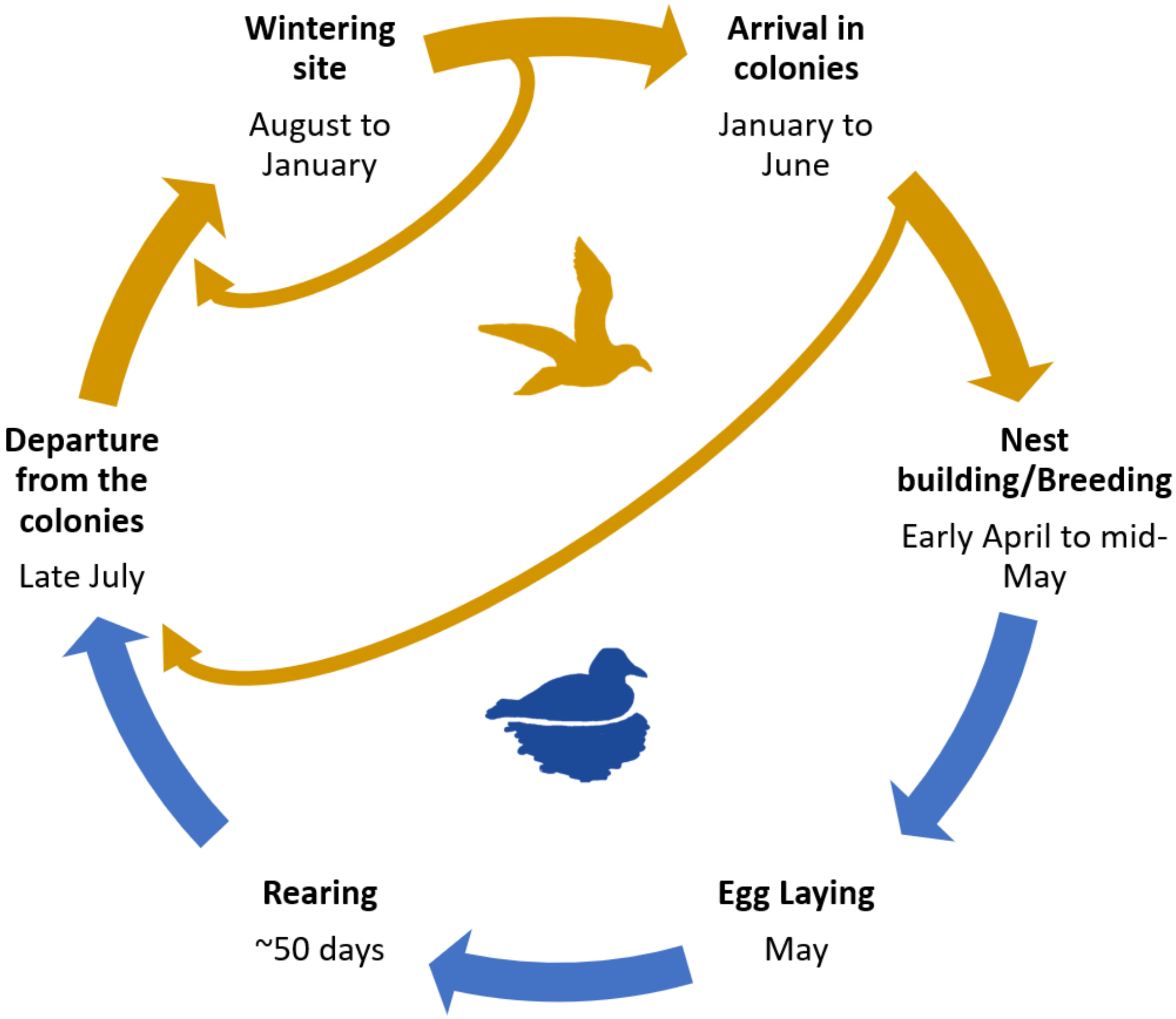

*Figure F1. Typical life-cycle of a black-legged kittiwake in the Cap-Sizun population with non-breeder state in yellow and breeder state in blue. Here, rearing corresponds to the total period of chick's dependency on its parents.*

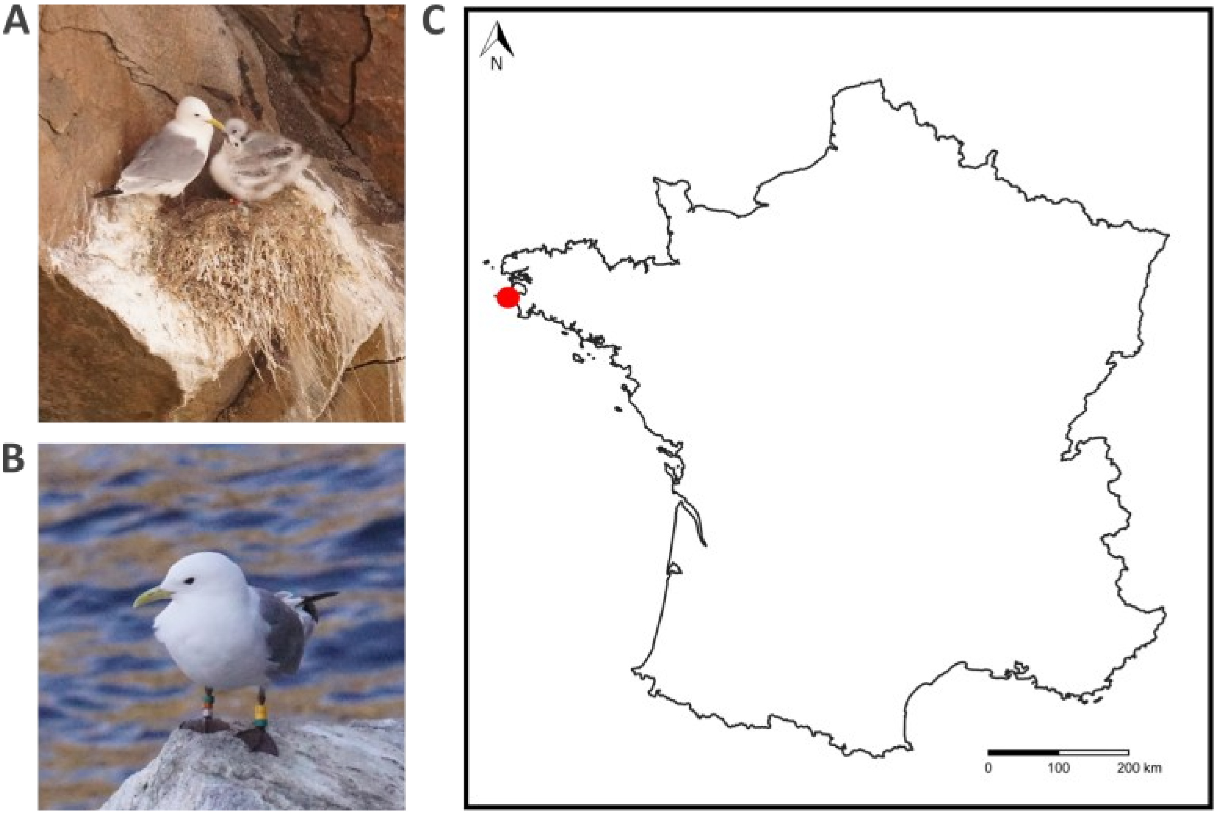

*Figure F2. Photos of newly ringed chicks on a completed nest with their parent (A) and a ringed adult (B) in the Kittiwake population in Cap Sizun, France (red dot in panel C). Here, birds are ringed with one metalic ring and 4 or 5 colored rings, following a protocol allowing to identify the individual and its cohort.*

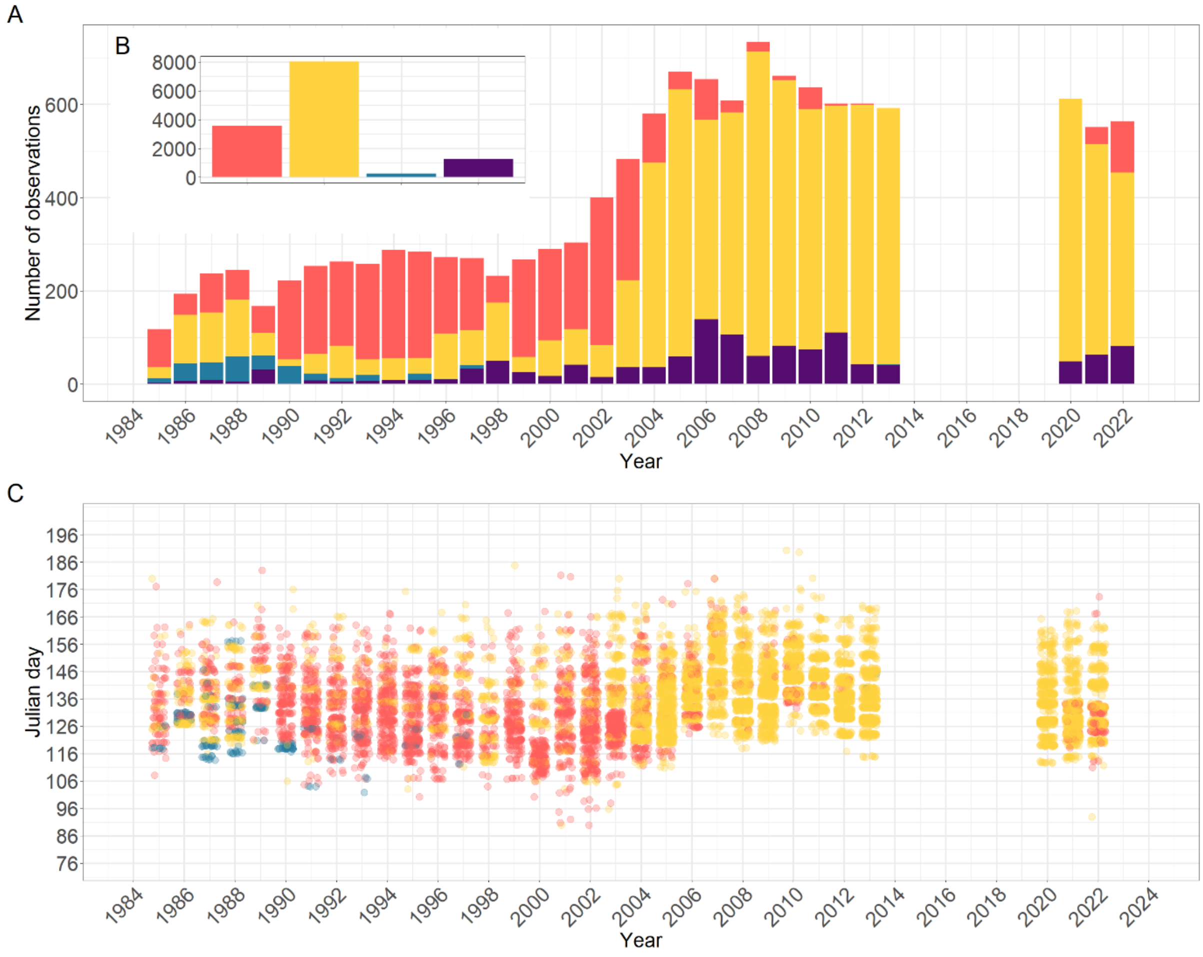

*Figure F3. Number of observations in (A) the overall dataset and (B) each year, and (C) annual date distribution of completed nest per year. Non-breeders are in purple and represent 9% of the overall observations. Other colours correspond to breeders with their associated precision on the nest completion date. Orange for a precision of 1 to 4 days (27% of the overall dataset), yellow for a precision of 5 to 12 days (61%), and blue for nests that were already completed on their first observation (3%).*

### Supplemental material G: Details of the environmental covariates and their biological effects

Here, we detail the construction of the two sets of environmental covariates implemented in our current illustrative analyses and the associated documented biological effects.

#### Defining seasonal progression and breeding season

The first set of covariates comprises two predictable photoperiod-related variables which non-tropical birds, such as kittiwakes, are widely known to use to physiologically perceive seasonal progression through the circadian clock (Ball & Balthazart, 2003; Jacobs & Wingfield, 2000). For breeding activities, photoperiod plays a key role in coordinating both gonad maturation and regression with the most appropriate period to breed (Ball & Balthazart, 2003; Dawson et al., 2001; Watanabe et al., 2007). First, when photoperiod matches the photoinducible phase of the circadian cycle, the circadian system synchronizes with the photoperiod signal, activating gonadal maturation and behaviors associated with breeding activities (e.g., territory defense, courtship, nest building, chick rearing). Therefore, gonadal growth is directly related to prevailing photoperiod (Ball, 1993; Ball & Balthazart, 2003; Dawson, 2015; Dawson et al., 2001). Second, gonadal regression implies photoperiodic time measurement, involving stored information of previously experienced light and dark cycle, allowing the organism to compare it with the present photoperiod to determine whether photoperiod is increasing or decreasing (Dawson, 2015; Dawson et al., 2001).

Consequently, we used two photoperiod-related variables in our analyses to represent the average response to the seasonal progression within a year in a constant environment: the daylength (in hours, denoted $\varphi$ ; Figure G1.A), to catch the physiological response to the increased photoperiod that leads to the start of the breeding season (Ball & Balthazart, 2003; Dawson, 2015; Dawson et al., 2001); and the difference in daylength between the focal and the previous day ($\Delta\varphi_d = \varphi_d - \varphi_{d-1}$, in hours; Figure G1.B), to approximate photoperiodic time measurement perception and the moment it will be physiologically too late for individuals to initiate breeding (Dawson, 2015; Dawson et al., 2001; Watanabe et al., 2007). The intrinsic relationships between these two variables and Julian day (measured in days since January 1st) are shown in Figure G1. $\varphi_d$ increases as the season progresses, while $\Delta\varphi_d$ have the opposite pattern, indicating that these two variables provide different information on the seasonal progression (i.e., prevailing photoperiod and increase or decrease of day length relative to previous day).

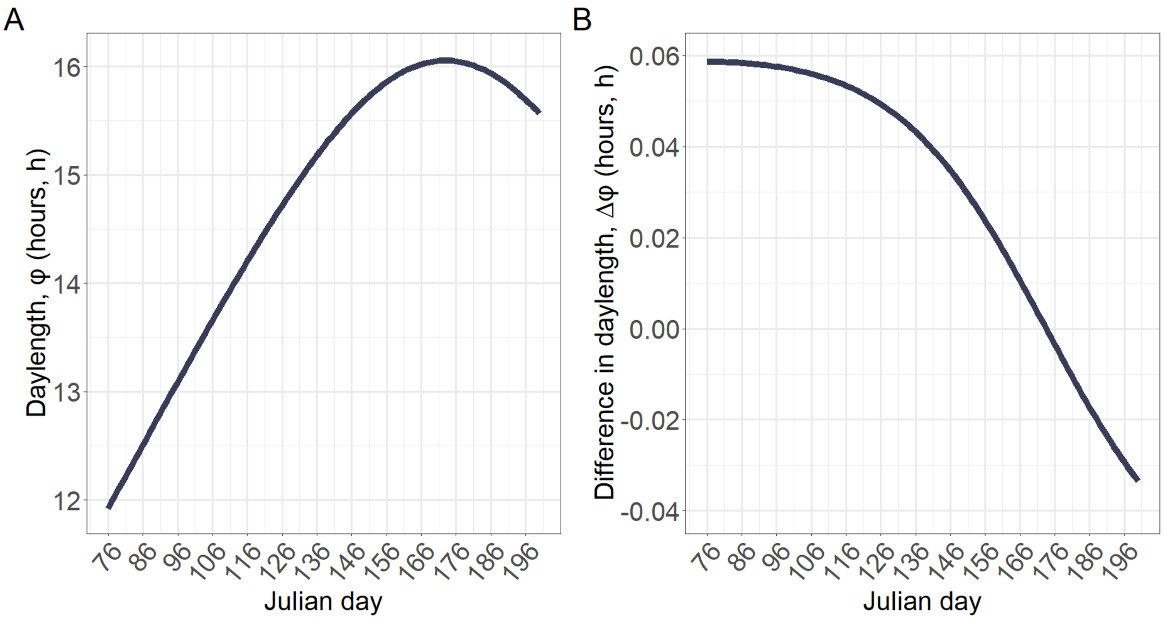

*Figure G1 : (A) Seasonal variation in daylength, $\varphi$ and (B) in the difference in daylength, $\Delta\varphi$, over the breeding season (from day 76 to 200).*

#### Temperature anomalies

The second set of environmental covariates aims to capture part of the stochastic component of the seasonal environment that varies among years.

For current illustrative purposes, we defined a temperature anomaly variable ($T$, °C) that quantifies within- and among-year variation in temperature. $T$ was computed in three steps (Figure G2). First, we extracted the raw daily mean air temperature over the duration of the study season and for each 32 years (1985-2013 and 2020-2022; data for 2014-2019 were not available) from the Météo-France online database (Météo-France, 2024) at the Pointe du Raz in Cap Sizun, located adjacent to the focal kittiwake colonies. Figure G2A shows that the mean air temperature increases as the season progresses and varies substantially among and within years. Second, we computed a rolling mean temperature over 20 days up to and including each focal day within D. Figure G2B shows that this resulting variable has the same among- and within-year trend as the mean air temperature but with reduced variance. Third, we computed the temperature anomaly variable. Figure G2C shows the relationship between annual variation of the temperature anomalies and the Julian day. $T$ fluctuates around zero over the duration of the study season without any particular trend with date. $T$ thereby enables capturing stochastic among- and within-year variation in temperature while avoiding collinearity with $\varphi$ and $\Delta\varphi$ (and hence, with the day).

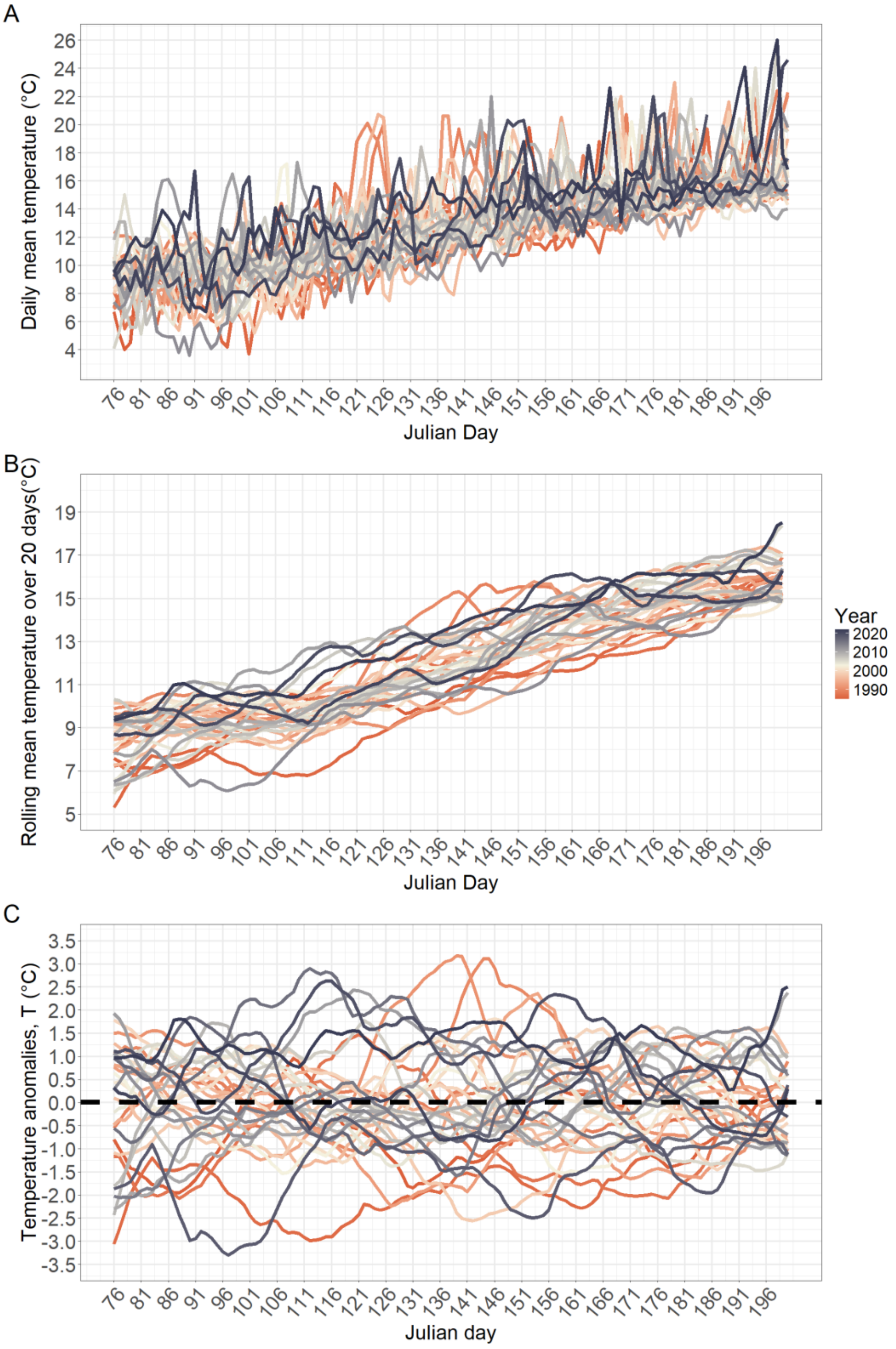

*Figure G2. Annual variation of (A) the mean daily temperature, (B) the daily rolled mean temperature over 20 days and (C) the temperature anomalies, T, over the duration of the study season (from day 76 to 200) for every 32 years (1985-2013 and 2020-2022) at the Pointe du Raz in Cap Sizun, which is the closest meteorological station to the colonies.*

### Supplemental material H: Kittiwake HM-PTM parameterization, priors and posterior distributions

Here, we detail the parameterization of the HM-PTM fitted to the kittiwake data (Equation 5 in the main text), information on model diagnostics, and present the priors and posterior distributions of the estimated liability reaction norm parameters (Figure H).

Figure H shows the posterior distributions of all LRN parameters (blue line) along with their associated prior distributions (red line). All posterior distributions peaked away from zero indicating clear evidence that the focal parameters were different from zero, except for the individual variance in the slope of temperature anomalies ($\sigma^2_{\beta_T}$) for which the mode was effectively on zero.

*Table H1: Priors for each of the LRN parameters.*

| **Parameters** | **Prior** |
|---|---|
| **Mean intercept ($\bar{\mu}$)** | Student-t(3, 0, 2.5) |
| **Mean slopes ($\bar{\boldsymbol{\beta}}_x$)** | Student-t(3, 0, 2.5) |
| **Variance in individual deviations from $\bar{\beta}_T$ ( $\sigma^2_{\beta_T}$)** | Half-Student-t(3, 0, 2.5) |
| **Variance in individual deviations from $\bar{\beta}_\varphi$ ( $\sigma^2_{\beta_\varphi}$)** | Half-Student-t(3, 0, 0.5) |
| **Variance in individual deviations from $\bar{\beta}_{\Delta\varphi}$ ( $\sigma^2_{\beta_{\Delta\varphi}}$)** | Half-Student-t(3, 0, 0.5) |
| **Variance in annual deviations from $\bar{\mu}$ ($\sigma^2_{\mu,y}$)** | Half-Student-t(3, 0, 2.5) |

*Table H2: Effective sample sizes (ESS) and $\hat{R}$ of the fitted HM-PTM for each of the LRN parameters.*

| Parameter | ESS | $\hat{R}$ |
|---|---|---|
| **Mean intercept ($\bar{\mu}$)** | 1,769 | 1.007 |
| **Mean slope on T ($\bar{\beta}_T$)** | 27,973 | 1.000 |
| **Mean slope on $\varphi$ ($\bar{\beta}_\varphi$)** | 3,124 | 1.002 |
| **Mean slope on $\Delta\varphi$ ($\bar{\beta}_{\Delta\varphi}$ )** | 2,082 | 1.003 |
| **Variance in individual deviations from $\bar{\beta}_T$ ( $\sigma^2_{\beta_T}$)** | 2,236 | 1.002 |
| **Variance in individual deviations from $\bar{\beta}_\varphi$ ( $\sigma^2_{\beta_\varphi}$)** | 495 | 1.011 |
| **Variance in individual deviations from $\bar{\beta}_{\Delta\varphi}$ ( $\sigma^2_{\beta_{\Delta\varphi}}$)** | 644 | 1.009 |
| **Variance in annual deviations from $\bar{\mu}$ ($\sigma^2_{\mu,y}$)** | 4,674 | 1.000 |

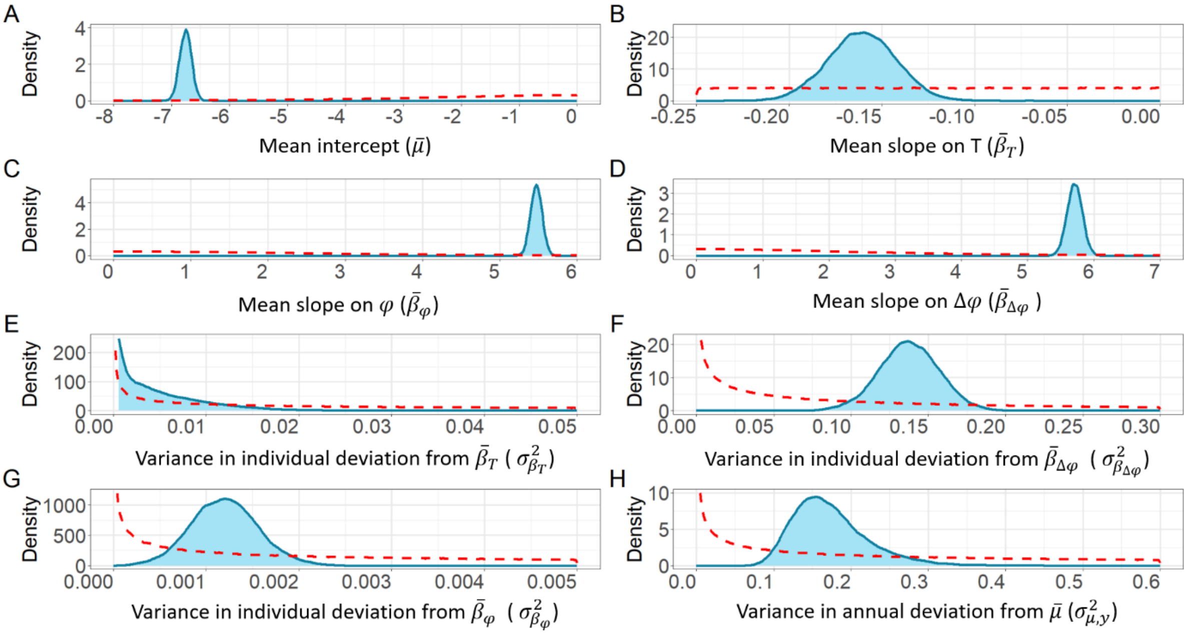

*Figure H. Posterior distributions of each of the parameters of the Latent Reaction Norm (panels A-H, blue) estimated by the model (Equation 5 in the main text) with their respective priors (red, details table H1). Panels show the posterior distribution and prior of (A) the mean intercept, (B) the mean slope on T, (C) the mean slope* on $\varphi$, *(D) mean slope on* $\Delta\varphi$, *(E) the variance in individual deviation from* $\bar{\beta}_T$, *(F) the variance in individual deviation from* $\bar{\beta}_\varphi$, *(G) the variance in individual deviation from* $\bar{\beta}_{\Delta\varphi}$, *and (H) the variance in annual deviation from* $\bar{\mu}$.

#### Supplemental material I: Defining the seasonal progression without the date

The main presented HM-PTM model structure (i.e. main text Equation 5) was chosen after testing a series of preliminary models in a developmental phase, which are summarised here. This illustrates how the (HM-)PTM approach can be used to test mechanistic liability-scale hypotheses (or conjectures), which would shape the phenological date distributions resulting from the LRN parameters.

For each hypothesis, we first devised a functional HM-PTM (i.e., achieving MCMC diagnostics indicating satisfactory convergence with largest $\hat{R} < 1.05$, Bulk and Tail ESS > 400; Vehtari et al., 2021). Second, we compared the predicted and observed annual distributions of nest completion dates to check whether the estimated LRN parameters generated biologically plausible phenological dates (Figure I). This allowed us to visually identify any non-trivial discrepancies that would reflect major lack of model fit exceeding the bias in raw observed nest completion dates stemming from the observation gaps.

More specifically, we devised a first HM-PTM (Equation I1) with just two of the three final environmental covariates: daily daylength, $\varphi_d$ (capturing the deterministic seasonal progression repeated each year) and temperature anomalies, $T_{d,y}$ (capturing stochastic interannual environmental variations). We achieved good MCMC diagnostics with the following formulation for the logit regression:

$$Logit\left(\Psi_{i,d,y}\right) = \bar{\mu} + \delta_{\mu,i} + \delta_{\mu,y} + \left(\bar{\beta}_T + \delta_{\beta_T,i}\right).T_{d,y} + \left(\bar{\beta}_\varphi + \delta_{\beta_\varphi,i}\right).\varphi_d, \quad \text{(I1)}$$

where all parameters are as defined in Equation 5 of the main text.

Here, the model-predicted seasonal trajectory of the population-wide mean daily nest completion probabilities ($\bar{\Psi}_{d,y}$) averaged across all years (i.e., green line on Figure I.A) broadly retrieved the expected bell shape through the seasonal progression. However, these predicted probabilities were relatively large at the beginning and end of the season where the true probabilities of nest completion should be virtually zero since kittiwakes were almost never recorded to complete nest building so early or so late (Figure I.A). Accordingly, while the mean nest completion dates in each year were adequately predicted, the predicted distributions were clearly too broad compared to the observed distributions (Figure I.B).

We hypothesized that these discrepancies could stem from the intrinsically non-linear relationship between $\varphi_d$ and date (Supplemental material G), which our initial model did not capture. We therefore fitted a second model that additionally included a quadratic effect of $\varphi_d$ with no among-individual variation (Equation I2):

$$Logit\left(\Psi_{i,d,y}\right) = \bar{\mu} + \delta_{\mu,y} + \left(\bar{\beta}_T + \delta_{\beta_T,i}\right).T_{d,y} + \left(\bar{\beta}_{\varphi} + \delta_{\beta_{\varphi},i}\right).\varphi_d + \bar{\beta}_{\varphi^2}.\varphi_d^2. \qquad \text{(I2)}$$

Here, we also removed the among-individual variance in the intercept. We did so because, given that $\varphi_d$ is fixed between years (instead of stochastically varying as the driving environmental variable in our simulations), $\sigma^2_{\mu,i}$ and $\sigma^2_{\beta_\varphi}$ are redundant parameters that cannot be properly estimated independently (as reflected by particularly bad model diagnostics, e.g. $\hat{R} >> 1.05$, Bulk and Tail ESS < 400, several transitions exceeding the maximum treedepth, and chains not mixing; Vehtari et al., 2021).

From this model, the seasonal trajectory of $\overline{\Psi}_{d,y}$ averaged across all years (i.e., yellow line on Figure I.A) was adequately estimated to be virtually zero in the beginning of the season. Accordingly, there was much better congruence of the left tails of predicted and observed date distributions (Figure I.C). However, discrepancies remained for the right tails of the distributions, with too many individuals predicted to complete their nest late compared to actual observations (Figure I.C).

Consequently, we tested a third alternative HM-PTM which, in line with wider understanding, assumes that kittiwake phenology is driven by the increase or decrease in daylength between consecutive days (Equation I3):

$$Logit\left(\Psi_{i,d,y}\right) = \bar{\mu} + \delta_{\mu,y} + \left(\bar{\beta}_T + \delta_{\beta_T,i}\right).T_{d,y} + \left(\bar{\beta}_{\Delta\varphi} + \delta_{\beta_{\Delta\varphi},i}\right).\Delta\varphi_d + \bar{\beta}_{\Delta\varphi^2}.\Delta\varphi_d^2, \qquad \text{(I3)}$$

Here, we replaced φ by $\Delta\varphi_d$, and retained the squared term. The results showed an opposite pattern from the previous model, where now the late seasonal trajectory of $\overline{\Psi}_{d,y}$ broadly captured the observed patterns (Figure I.A) but $\overline{\Psi}_{d,y}$ was too high early in the season. Accordingly, the date distributions indicated that the model predicted far too many individuals to complete their nest early compared to the raw observations (Figure I.D).

Altogether, these exploratory analyses indicated that both φ and Δφ can act as complementary covariates that combine to shape the seasonal distribution of daily probability to initiate breeding and resulting predicted annual dates distributions, where φ informs on the duration of the current day and Δφ informs on whether the focal day is longer or shorter than the previous one. Therefore, to better catch the underlying mechanisms that leads to the phenological date, we included both covariates in our final HM-PTM.

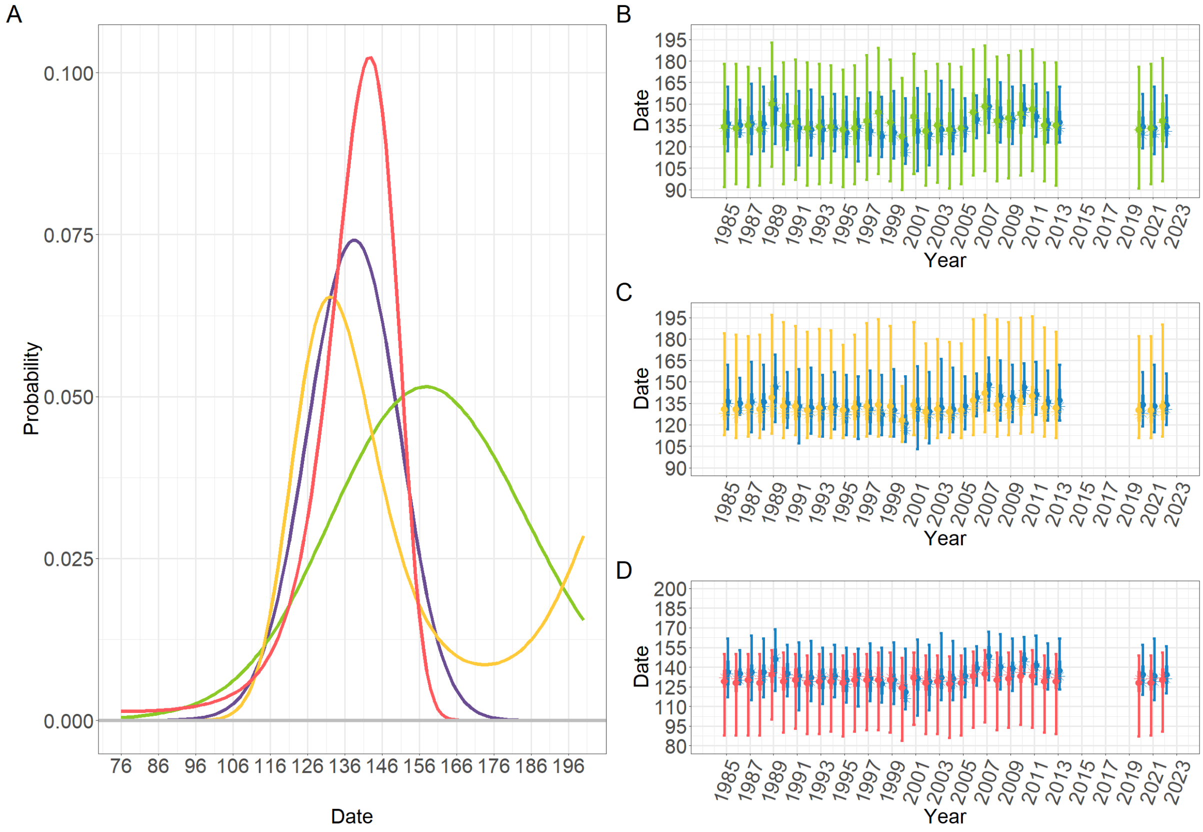

*Figure I. (A) Seasonal trajectory of the predicted daily probability of nest completion averaged across all years and (B, C, D) the resulting observed (blue) and predicted (other colours) annual date distribution for each model tested. Predictions by the first model (Equation I1) are represented in green (A and B); the model with the quadratic effect on φ (Equation I2) are in yellow (A and C); the model with Δφ (Equation I3) are in pink (A and D). On panel A, the purple curve is from the model presented in the main manuscript. Dots and stars on the date distributions on graphs B, C and D represent respectively the annual mean and median dates of nest completion. Note that the predicted annual distributions are part of preliminary analyses where the method to infer predicted dates were slightly different than the one detailed in the main text. Here, phenological date distribution have been inferred by simulating a large number of liability reaction norms using point estimates of the posterior median of each estimated liability scale parameters. Although, the resulting annual predicted date distributions are similar with both methods, the method used in the main article enables propagating estimation uncertainty in the predicted date distributions.*

#### Supplemental material J: Relationship between the resighting regime and the predicted annual date distributions

In this section, we compare the nest completion dates predicted by the HM-PTM analysed in the main text with the raw observed dates, given the data gaps resulting from the field observation regime (detailed Supplemental material F). We further evaluate the capacity of the HM-PTM by comparing the observed and predicted date distributions resulting from our simulated dataset with a balanced design, with and without gaps (detailed in main text and Supplemental material E).

Using our main HM-PTM fitted to the kittiwake data (main text Equation 5), the means of the predicted annual distributions of nest completion dates are typically earlier than the observed means by 5 days on average (range 13 days earlier to 2 days later; Figure 5 of the main text). The 2.5% quantile is typically earlier by 9 days (maximum 23 days earlier, minimum 7 days later) and the 97.5% quantile is typically earlier by 7 days (maximum 16 days earlier, minimum 0 days later). The pattern of among-year variation in magnitudes of these discrepancies (Figure 5) coincided with the change of resighting regime (presented in Supplemental material F): 1 to 4 days between successive observations before 2004 and 5 to 12 days from 2004. Overall, this indicates that differences between observed and predicted date distributions may well be explained by bias in the date distributions directly derived from the raw data due to incompleteness of field observations.

Further, using our simulation dataset, we compared the annual date distributions simulated before and after implementation of 5-day gaps versus the annual date distributions predicted by the HM-PTM (Figure J) fitted with the dataset with 5-day gaps (details on the dataset and the models presented section “Model Evaluation” and Figure 3 in the main text, and Supplemental material E). For the two types of simulated environments (i.e., linear and non-linear) and the two types of Latent Reaction Norm (LRN) simulated, the annual date distribution predicted by our model is the same as the one simulated before implementation of the resighting gaps (i.e., the "real" dates). This clearly demonstrates the capacity of the (HM-)PTM to accurately predict the true distributions of event dates by correcting for the bias in the raw observations stemming from incompleteness in observations of individual state sequences. Thereby, the results in Figure J strongly support the point that the discrepancies between observed and predicted date distribution of kittiwake nest completion could be largely reflecting the bias due to incomplete field observations.

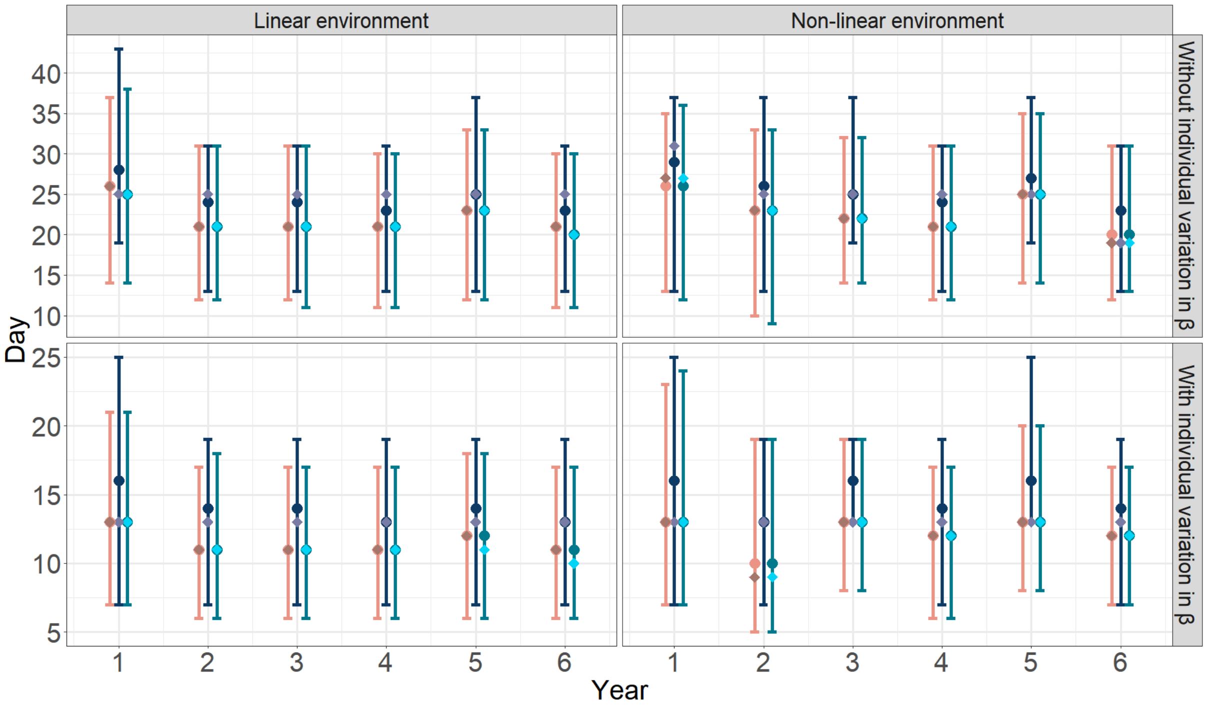

*Figure J. Summarised annual distributions of simulated (blue) and predicted (pink) dates for the 2 simulated environment (columns) and the two types of simulated Latent Reaction Norm (LRN; rows). Light blue distributions represent the simulated dates under a balanced design without observations gaps (i.e., simulated "real" dates). Dark blue distributions are the simulated dates resulting from the same dataset after applying a 5-day gap observational regime. Pink corresponds to the annual date distributions predicted by the Hidden Markov-Phenological Threshold Model (HM-PTM) presented in dark blue in figure 3 in the main text. The dots, diamond and bars represent respectively the means, medians, and 95% of the annual date distributions.*
